# Integrated culturing, modeling and transcriptomics uncovers complex interactions and emergent behavior in a synthetic gut community

**DOI:** 10.1101/299644

**Authors:** Kevin D’hoe, Stefan Vet, Karoline Faust, Frédéric Moens, Gwen Falony, Didier Gonze, Verónica Lloréns-Rico, Lendert Gelens, Jan Danckaert, Luc De Vuyst, Jeroen Raes

## Abstract

Whereas the composition of the human gut microbiome is relatively well resolved, predictive understanding of its response to perturbations such as diet shifts is still lacking. Here, we followed a bottom-up strategy to explore human gut community dynamics. We established a synthetic community composed of three representative human gut isolates in well-controlled conditions *in vitro.* We then explored species interactions by performing all mono- and pair-wise fermentation experiments and quantified with a mechanistic community model how well tri-culture dynamics was predicted from mono-culture data. With the model as a reference, we demonstrated that species grown in co-culture behaved differently than in mono-culture and confirmed their altered behavior at the transcriptional level. In addition, we showed with replicate tri-cultures and in simulations that dominance in tri-culture sensitively depends on initial conditions. Our work has important implications for gut microbial community modeling as well as ecological interaction detection from batch cultures.

## Introduction

The human gut microbiome is a complex, spatially heterogeneous and dynamic system consisting of hundreds of species interacting with each other and with the human host. It is a daunting task to develop predictive models for such a system, yet the potential rewards are high and would, for instance, enable targeted interventions to shift dysbiotic communities towards more healthy states. Two conditions need to be fulfilled for predictive models to be successful: first, the system has to be sufficiently well characterized to build the model and second, the dynamics should be generally deterministic. First successes in modeling the behavior of gut microbial communities give reason for cautious hope (Buffie, Bucci et al., 2015, Cremer, Arnoldini et al., 2017, Muñoz-Tamayo, Giger-Reverdin et al., 2016, Stein, Bucci et al., 2013). Most of these studies took a top down approach, where the change in composition of an entire community *in vivo* is modeled. For instance, Cremer and colleagues predicted the ratio of Firmicutes and Bacteroidetes in fecal samples as a function of estimated water content and nutrient influx using a diffusion model (Cremer et al., 2017). Others have fitted population models to time series of taxon (mostly genus) abundances obtained from 16S rRNA gene sequencing. For instance, one study fitted a variant of the generalized Lotka-Volterra (gLV) model to a cecal gut time series of mice exposed to a pathogen (*Clostridium difficile*), an antibiotic or both, thereby inferring the interactions between different genera (Stein et al., 2013). The time series predicted with the parameterized gLV model agreed well with observations. The same approach was also used to predict species that inhibit *C. difficile* growth in murine and human microbiota, one of which significantly lowered mortality when transferred to mice before infection with *C. difficile* (Buffie et al., 2015).

Despite these successes, the gLV model and its variants have several drawbacks that limit their widespread application. Importantly, they assume that community dynamics can be predicted from pair-wise interactions and that the interaction mechanism can be ignored. These assumptions have recently been challenged both experimentally and computationally: Friedman and colleagues experimentally quantified the accuracy reached when predicting the behavior of more complex soil communities from species pairs (Friedman, Higgins et al., 2017), whereas Momeni and colleagues systematically compared gLV models of metabolite-mediated species interactions to their mechanistic counterparts *in silico* (Momeni, Xie et al., 2017). While the authors in the former case concluded that the behavior of larger communities could to a considerable extent be predicted from that of smaller ones, the latter showed that the (extended) gLV model cannot describe accurately several common types of interaction mechanisms.

An alternative to the gLV model and its variants are mechanistic models, which account for nutrient-mediated interactions by explicitly describing the dynamics of the produced and consumed compounds (see (Momeni et al., 2017) and references therein). They thus require more system knowledge than generic gLV and related models do. However, most members of the gut community have not been thoroughly characterized, and little is known about their responses to different nutrients, pH values and interaction partners - even for those that have been studied more closely. It is challenging to obtain this type of biological knowledge and to resolve interaction mechanisms *in vivo.* However, *in vitro* studies not only allow acquiring detailed knowledge of the microorganisms’ pH and nutrient preferences, but also of their behavior in the presence of other microorganisms. *In vitro* studies of human gut microorganisms have a long tradition and have been carried out in several manners. Classical mono- and co-culture studies in batch and chemostat fermentors have explored nutrient preferences and interaction mechanisms (Falony, Calmeyn et al., 2009a, Falony, Vlachou et al., 2006, Moens, Verce et al., 2017, Moens, Weckx et al., 2016, Rivière, Selak et al., 2016). Artificial gut systems, such as the TNO In Vitro Model of the Colon (TIM-2) (Venema, 2015) and the Simulation of the Human Intestinal Microbial Ecosystem (SHIME) (T. Van de Wiele, P. Van den Abbeele et al., 2015), seek to reproduce as closely as possible the conditions of the human gastro-intestinal tract in a well-controlled manner. They are composed of several compartments connected with peristaltic pumps that vary in their pH to mimic the pH gradient in the human intestines. In addition, the SHIME system can be extended with a module dedicated to the investigation of human-microbiome interactions (HMI) (Marzorati, Vanhoecke et al., 2014). The gut community has also been studied at smaller scales, in minibioreactor arrays (Auchtung, Robinson et al., 2015) and with gut-on-chip microfluidic devices (Kim, Huh et al., 2012, Shah, Fritz et al., 2016). However, in most cases, gut simulators are inoculated with fecal material. In the range from top-down to bottom-up approaches of gut microbial community dynamics, these can be considered as intermediate cases, where the host is eliminated, but the community is not further simplified. The goal of these studies is usually to quantify the behavior of the entire community under different conditions. In the case of HuMiX and SHIME’s HMI module, the interaction of particular gut microorganisms with epithelial cells is targeted (Marzorati et al., 2014, Shah et al., 2016). Since the exact composition of fecal material (which also includes bacteriophages and fungi) is difficult to resolve, it is hard to track each member in such a community. Although the *in vitro* dynamics of colon (Kettle, Louis et al., 2015) and rumen (Muñoz-Tamayo et al., 2016) communities has been described with mechanistic models previously, these models did not account for the behavior at species level, but grouped species with similar metabolic activities into guilds. Whereas it is of interest to model guild dynamics, the resolution of guild-level models may be insufficient to understand microbial community dynamics in the gut. Species in the same guild do not necessarily respond in the same manner to altered environments and perturbations. Guild definitions are arbitrary to an extent, and gut bacteria with flexible metabolic strategies may change their guild membership. In addition, the concepts of tipping elements (Lahti, Salojãrvi et al., 2014) and strongly interacting species (Gibson, Bashan et al., 2016) suggest that particular species can have a disproportionate impact on gut community dynamics.

In our opinion, experiments with defined communities of known composition, grown in well-controlled conditions, are crucial to learn more about the interactions of gut species and how these shape community dynamics. Well-controlled *in vitro* experiments are also necessary for the development and validation of predictive models of gut microbial communities. However, only few *in vitro* experiments with defined gut communities have been reported to date (Newton, Macfarlane et al., 2013, Trosvik, Rudi et al., 2008, Trosvik, Rudi et al., 2010) and none have to our knowledge employed mechanistic models to predict community dynamics.

The objective of the present study was to establish a defined gut community composed of representative human gut strains, to study their interactions under well-controlled conditions *in vitro* and to validate a quantitative mechanistic model by predicting community behavior in a tri-culture with parameters from mono-culture data. Whereas mechanistic models have been tested in this manner before for a cystic fibrosis community (Schmidt, Riedele et al., 2011), such an approach has not yet been applied to a synthetic gut community.

To reach our objective, we established three gut bacterial strains, described below, in an optimized mMCB medium (Moens et al., 2016) in 2-L laboratory fermentors run in batch mode. To measure kinetic parameters, we carried out mono-cultures and then cultivated all three strains together in a tri-culture. In addition, we quantified the dynamics of each of the three possible pair-wise combinations and performed all experiments in at least two biological replicates. For each cultivation experiment, we took samples at 15 to 18 time points over two days. For all samples, we quantified optical density, counted cells with flow cytometry, measured the concentration of fructose and short-chain fatty acids (formate, acetate, propionate, butyrate and lactate) and quantified bacterial abundance with TaqMan qPCR using species-specific primers and probes. In addition, we quantified the concentration of CO_2_ and H_2_ in the headspace during fermentation in a semi-continuous fashion. Finally, we sequenced the total RNA in selected samples. Figure 1 summarizes our approach. To our knowledge, this is the first study that investigates a synthetic gut community with a combination of mono- and co-cultures, mechanistic modeling and gene expression analysis.

**Figure 1.**
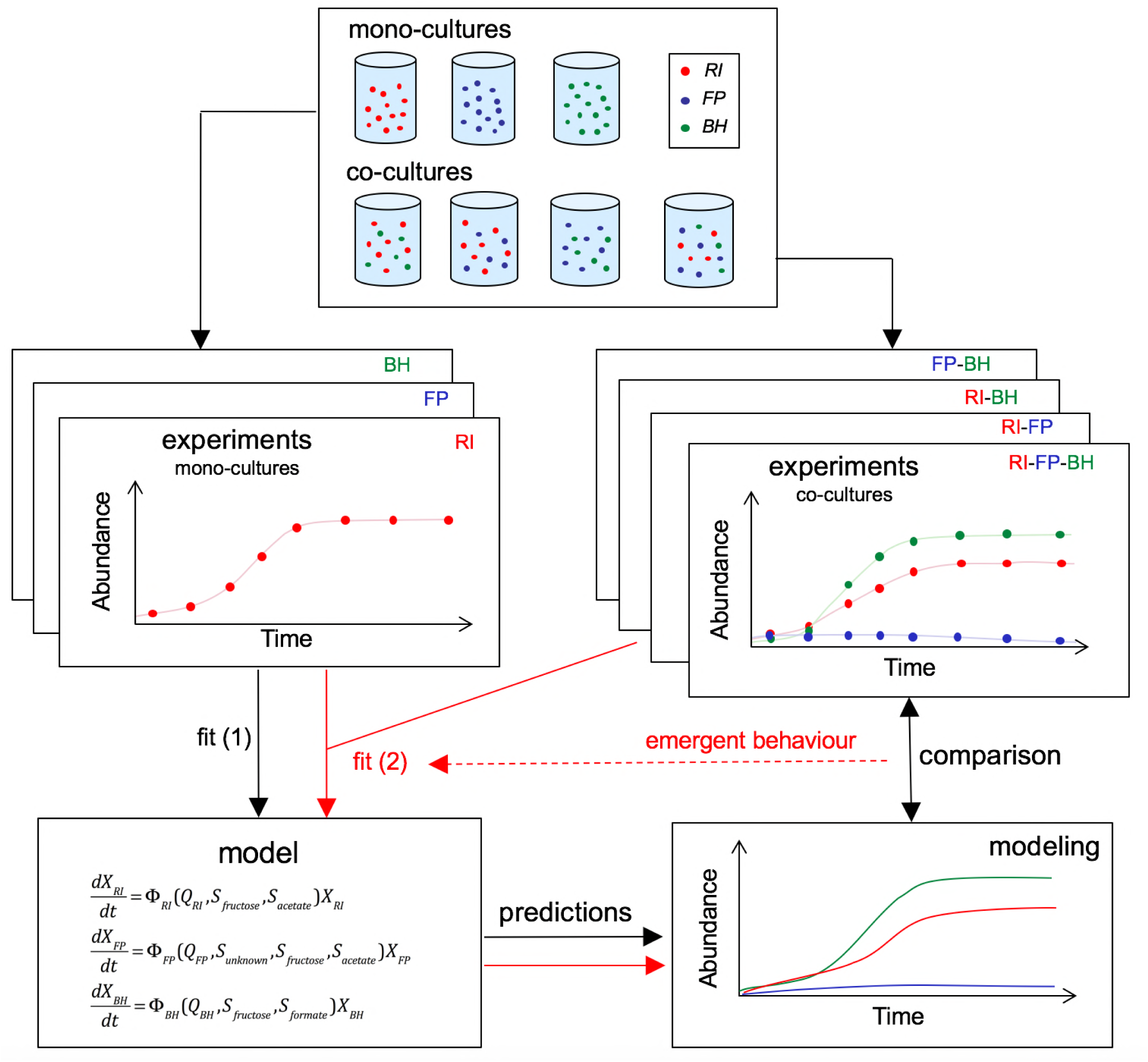
Scheme summarizing the experimental set-up and modeling approach. A mechanistic model of a three-strain community consisting of *Roseburia intestinalis* (RI), *Faecalibacterium prausnitzii* (FP) and *Blautia hydrogenotrophica*(BH) is parameterized on mono-cultures, but does not describe tri-culture dynamics well. Data from co-cultures are taken into account to improve the goodness of fit to the tri-culture data, thereby indicating emergent behavior.

Our synthetic community was composed of *Faecalibacterium prausnitzii* A2-165 (Duncan, Hold et al., 2002b) (FP), *Roseburia intestinalis* L1-82 (Duncan, Hold et al., 2002a) (RI) and *Blautia hydrogenotrophica* S5a33 (Bernalier, Willems et al., 1996) (BH), all three of which were isolated from human feces. We carefully selected our community members according to several criteria. First, we targeted abundant and typical members of the human gut microbiome. According to the Flemish Gut Flora Project (FGFP), for which fecal samples of 1,106 healthy Flemish adults were sequenced, *Faecalibacterium* is the top-abundant genus and a *Faecalibacterium* OTU the top-abundant OTU in the healthy gut (Falony*, Joossens* et al., 2016). *Roseburia* and *Blautia* are both among the top 10 most abundant gut genera. In addition, *Faecalibacterium* and *Roseburia* OTUs were present in all sequenced samples, whereas Blautia OTUs were present in 99.9% of the samples. The selected *Faecalibacterium* and *Roseburia* strains were also among the strains with more than 1% abundance in several of the whole-genome-shotgun sequenced stool samples from the Human Microbiome Project (Kraal, Abubucker et al., 2014). Thus, our community is composed of representative human gut strains.

Second, we specifically targeted butyrate producers (RI and FP). Butyrate is an important energy source for gut epithelial cells and is beneficial for gut health (Geirnaert, Calatayud et al., 2016, Rivière et al., 2016). Butyrate producers are often found to be enriched in healthy as compared to dysbiotic gut microbiota (Antharam, Li et al., 2013, Rivera-Chávez, Zhang et al., 2016). Thus, high butyrate production will likely be a quality criterion for bacterial cocktails designed for therapeutic purposes. Our synthetic community allows studying more closely the factors that influence butyrate production.

Third, we designed our community with a large number of potential interactions. As Figure 2 illustrates, our community contains multiple cross-feeding and competitive interactions. For instance, FP needs acetate, which is produced by BH. Vice versa, BH grows on the formate generated by FP. Both cross-feeding relationships together constitute a mutualistic interaction. In parallel, both species compete for the energy source, fructose. The same relationships also occur between RI and BH (which can exchange hydrogen and CO_2_ in addition to formate), while RI and FP compete for fructose and acetate. This system also constitutes a rare example of two species pairs that simultaneously compete and mutually cross-feed.

**Figure 2:**
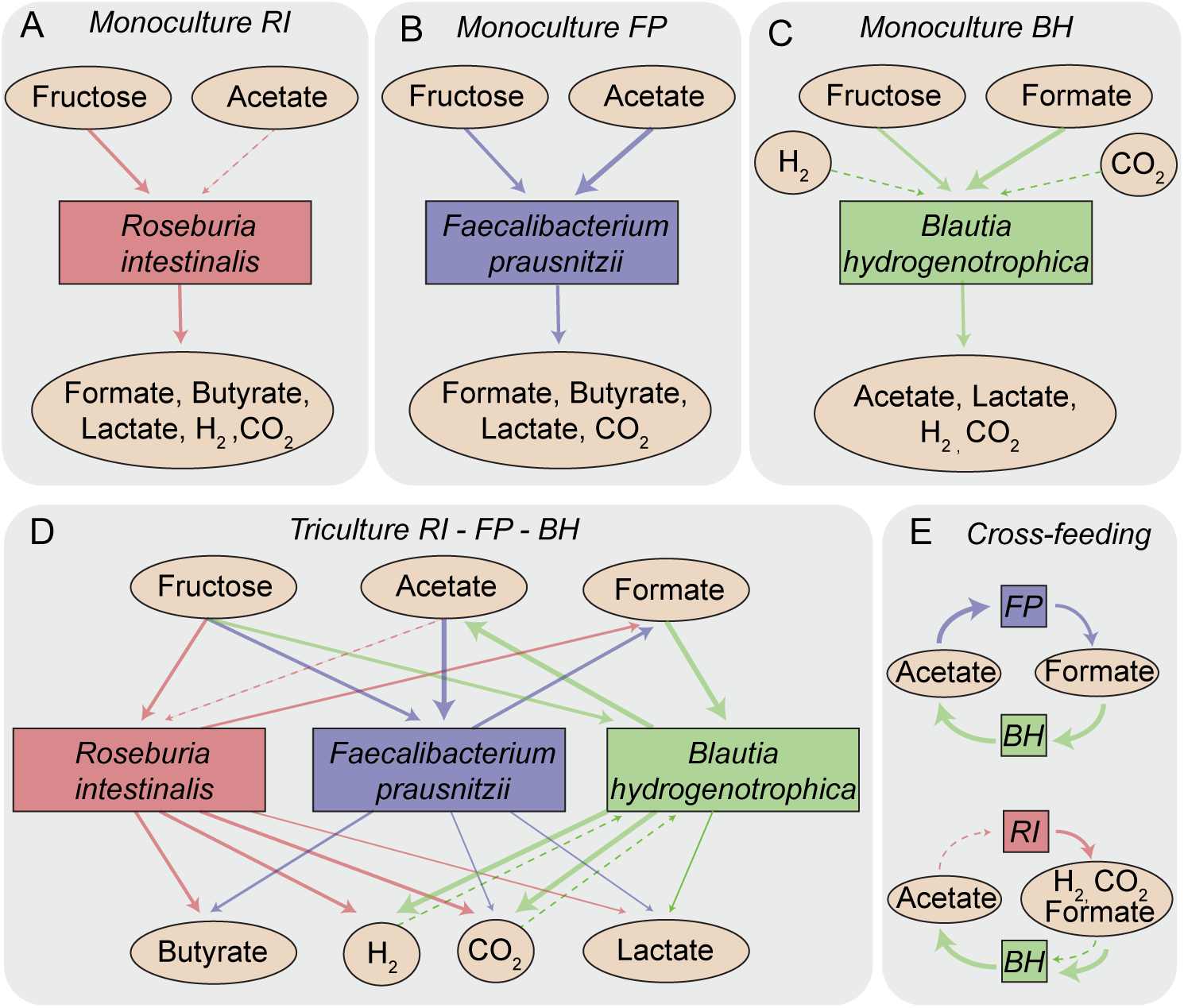
Overview of metabolite-mediated strain interactions. (A-C) Strain-specific metabolite consumption and production. (D) Metabolite-mediated interactions present in the tri-culture. (E) Cross-feeding interactions between *Faecalibacterium prausnitzii* (FP) and *Blautia hydrogenotrophica* (BH) as well as *Roseburia intestinalis* (RI) and BH. The dashed arrow from acetate to *Roseburia intestinalis* denotes net acetate consumption. The dashed arrows from hydrogen and CO_2_ to *Blautia hydrogenotrophica* indicate the potential of this bacterium to grow autotrophically on these gasses.

Finally, we only considered human gut species for which at least a draft genome was available, to ease primer design and the interpretation of RNA-seq data.

## Results

### *Blautia hydrogenotrophica* is metabolically versatile

We first confirmed the cross-feeding interactions postulated for BH with small-volume screening experiments, where the pH was not kept constant and the atmosphere contained 10% CO_2_ and 10% hydrogen gas. We found that under these conditions, BH was able to grow heterotrophically on formate and autotrophically as described previously on CO_2_ and hydrogen gas (Bernalier et al., 1996), presumably in both cases via the Wood-Ljungdahl pathway, of which all genes were located in BH’s genome (Rey, Faith et al., 2010).

We also found in agreement with (Rey et al., 2010) that BH grew on fructose, oligofructose and glucose, albeit more slowly than on formate. In agreement with (Bernalier et al., 1996), we detected lactate in addition to acetate for these substrates. We thereby confirmed the potential for fructose competition between BH on the one hand and RI or FP on the other in our medium. BH also consumed small concentrations of galactose, but did not consume fucose, inulin or lactate. For BH grown on fructose, we confirmed slow growth and lactate production in addition to CO_2_, H_2_ and acetate in the fermenter. Notably, when growing BH on formate in the fermenter, carbon dioxide and hydrogen gas were produced besides acetate, but no lactate.

In conclusion, our work showed that *Blautia hydrogenotrophica* is a surprisingly versatile member of the human gut ecosystem.

### Mono-culture dynamics does not follow standard kinetics

We employed pH-controlled monocultures to characterize the properties and growth kinetics of the individual strains in our model. Table 1 provides an overview for all fermentation experiments carried out, whereas Supplementary Table 1 gives additional information for each experiment.

**Table 1:** Overview of fermentation experiments and model fitting results (RMSE: root mean square error). Mean and standard deviations across biological replicates are reported.

| Strains | Number of biological replicates | Added energy source and/or co-substrate | Mean consumption/production ± standard deviation (mM) of metabolites |  |  |  |  |  |  | Carbon recovery (%) | Selected for parameterization 1 | RMSE parameterization 1 | Selected for parameterization 2 | RMSE parameterization 2 |
| --- | --- | --- | --- | --- | --- | --- | --- | --- | --- | --- | --- | --- | --- | --- |
|  |  |  | Fructose | Acetate | Butyrate | Formate | Lactate | H<sub>2</sub> | CO<sub>2</sub> |  |  |  |  |  |
| Monoculture fermentations |  |  |  |  |  |  |  |  |  |  |  |  |  |  |
| *R. intestinalis* DSM 14610<sup>T</sup> | 4 | Fructose/ Acetate | -48.8 ± 0.9 | -20.6 ± 2.4 | 58.5 ± 3.9 | 6.8 ± 1.2 | 3.9 ± 1.1 | 72.2 ± 4.9 | 84.0 ± 0.8 | 98.3 ± 6.5 | Yes | 0.17 ± 0.04 | No | 0.83 ± 0.13 |
| *F. prausnitzii* DSM 17677<sup>T</sup> | 3 | Fructose/ Acetate | -23.1 ± 1.1 | -13.5 ± 0.8 | 29.8 ± 0.2 | 22.7 ± 0.9 | 1.8 ± 0.2 | 0.2 ± 0.2 | 19.3 ± 1.6 | 100.9 ± 3.4 | Yes | 0.15 ± 0.08 | Yes | 0.15 ± 0.08 |
| *B. hydrogenotrophica* DSM 10507<sup>T</sup> | 3 | Fructose/ Formate | -19.0 ± 5.6 | 23.0 ± 6.6 | 0.0 ± 0.0 | -36.1 ± 1.8 | 6.0 ± 1.2 | 31.5 ± 7.7 | 26.0 ± 5.3 | 60.0 ± 2.1 | Yes | 1.43 ± 0.27 | No | 3.53 ± 1.04 |
| Bi-culture fermentations |  |  |  |  |  |  |  |  |  |  |  |  |  |  |
| *R. intestinalis* DSM 14610<sup>T</sup>/ *F. prausnitzii* DSM 17677<sup>T</sup> | 2 | Fructose/ Acetate | -47.7 ± 2.2 | -18.1 ± 2.7 | 56.3 ± 4.5 | 7.6 ± 2.7 | 4.3 ± 3.3 | 114 ± 16 | 85.7 ± 4.1 | 102.8 ± 0.3 | No | 0.77 ± 0.06 | No | 0.68 ± 0.03 |
| *R. intestinalis* DSM 14610<sup>T</sup>/ *B. hydrogenotrophica* DSM 10507<sup>T</sup> | 2 | Fructose/ Acetate | -46.5 ± 1.5 | -7.5 ± 1.9 | 53.5 ± 4.7 | -0.5 ± 0.8 | 2.2 ± 0.3 | 53.0 ± 3.0 | 74.9 ± 4.1 | 100.2 ± 0.6 | No | 0.78 ± 0.36 | Yes | 0.3 ± 0.04 |
| *R. intestinalis* DSM 14610<sup>T</sup>/ *B. hydrogenotrophica* DSM 10507<sup>T</sup> | 2 | Fructose | -48.0 ± 0.2 | 44.9 ± 5.4 | 27.9 ± 3.0 | -1.0 ± 0.0 | 5.4 ± 2.4 | 33.1 ± 3.2 | 47.4 ± 3.9 | 91.6 ± 1.1 | No | 0.9 ± 0.17 | No | 0.32 ± 0.04 |
| *F. prausnitzii* DSM 17677<sup>T</sup>/ *B. hydrogenotrophica* DSM 10507<sup>T</sup> | 2 | Fructose/ Acetate | -49.3 ± 1.0 | 41.4 ± 11.7 | 30.7 ± 1.8 | -1.2 ± 0.0 | 3.9 ± 1.6 | 56.2 ± 37.1 | 54.8 ± 33.3 | 91.6 ± 9.4 | No | 0.6 ± 0.14 | Yes | 0.26 ± 0.13 |
| *F. prausnitzii* DSM 17677<sup>T</sup>/ *B. hydrogenotrophica* DSM 10507<sup>T</sup> | 1 | Fructose | -47.0 | 62.5 | 25.5 | -1.1 | 4.5 | 62.9 | 63.6 | 107.4 | No | 0.63 | No | 0.46 |
| Tri-culture fermentations |  |  |  |  |  |  |  |  |  |  |  |  |  |  |
| *R. intestinalis* DSM 14610<sup>T</sup>/ *F. prausnitzii* DSM 17677<sup>T</sup>/ *B. hydrogenotrophica* DSM 10507<sup>T</sup> | 6 | Fructose | -48.9 ± 1.3 | 38.4 ± 16.9 | 32.5 ± 5.5 | -1.2 ± 0.1 | 7.4 ± 1.7 | 61.9 ± 3.2 | 57.1 ± 9.2 | 97.0 ± 3.1 | No | 0.78 ± 0.27 | No | 0.58 ± 0.11 |

RI consumed fructose and produced butyrate, CO_2_ and hydrogen gas, as described previously (Falony, Verschaeren et al., 2009c) as well as small amounts of lactate and formate (Figure 3A). Interestingly, there was no net consumption of acetate when more fructose than acetate was provided. While net acetate consumption has been found to correlate negatively with hydrogen production (Falony et al., 2009c), we saw here that it also depends on the ratio of initial fructose and acetate. When given in equal concentrations, RI partially consumed acetate. In all further experiments, when adding acetate, it was added at the same concentration as fructose.

**Figure 3:**
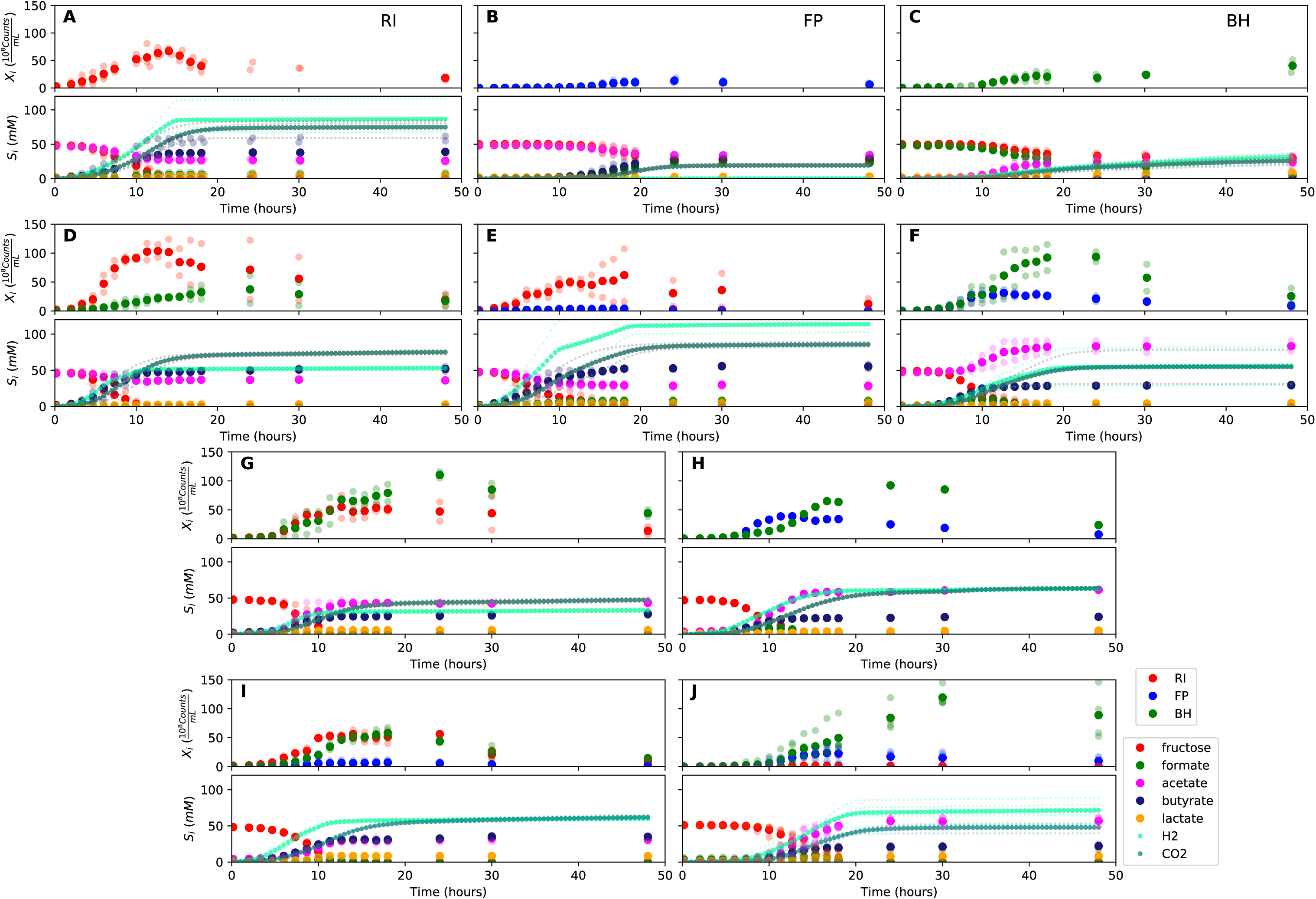
Summary of fermentation data. Biological replicates are plotted together in one panel, with their mean shown in bold. For each set of experiments, species abundances quantified by qPCR are plotted in the top half of the panel and metabolite concentrations in the bottom half. (A-C) Mono-cultures of *Roseburia intestinalis* (RI), *Faecalibacterium prausnitzii* (FP) and *Blautia hydrogenotrophica* (BH). (D-F) The three co-culture combinations of RI, FP and BH with initial acetate. (G-H) Co-cultures of RI versus BH and FP versus BH without initial acetate. (I-J) The tri-culture replicates are separated into those dominated by RI and BH (I) and those dominated by FP and BH (J).

FP in mono-culture produced formate, less CO_2_ and butyrate than RI and no hydrogen gas, but did not entirely consume fructose (Figure 3B). After having excluded exposure to oxygen (by adding oxygen gas via sterile water), redox potential (by continuously adding the oxidizing agent potassium ferrocyanide trihydrate), pH (lowered to 5.8), a threshold requirement for fructose (halving the fructose concentration did not stop its consumption) or end-product inhibition (by adding initial butyrate) as explanations, we found that doubling the concentration of yeast extract lowered fructose concentrations. Since adding fresh but autoclaved medium during the fermentation did not lower fructose concentrations, we assume that FP was growth-limited by (a) heat-labile co-factor(s) present in the yeast extract. A recent flux balance analysis with a manually curated metabolic reconstruction suggests that FP requires several amino acids (L-alanine, L-cysteine, L-methionine, L-serine and L-tryptophan) and the co-factors biotin (vitamin B_7_), cobalamin (vitamin B_12_), folic acid (vitamin B_9_), hemin, nicotinic acid, pantothenic acid and riboflavin (vitamin B_2_) for growth (Heinken, Khan et al., 2014). With the exceptions of cobalamin and externally supplied hemin, these nutrients should be present in yeast extract according to the metabolic reconstruction of *Saccharomyces cerevisiae* iMM904 (Mo, Palsson et al., 2009) and the amino acids should furthermore be present in other medium components (bacteriological peptone, soy peptone and tryptone). Heinken and colleagues also predicted that FP can grow in the presence of oxygen, which is in agreement with our observation that addition of low concentrations of oxygen does not alter FP’s growth curve (Heinken et al., 2014). BH produced acetate, hydrogen, CO_2_ and small concentrations of lactate, while consuming formate almost entirely (Figure 3C). It also consumed fructose, but did not deplete it. Whereas the carbon recovery for the other mono-cultures was close to 100%, it only reached 60% for BH in mono-culture on formate and fructose.

These unexpected behaviors defy simple kinetic models and necessitate further adjustment of the equations.

### Prediction accuracy of model parameterized on mono-cultures is species-dependent

We designed a model that described the dynamics of each species and of key compounds (including fructose, formate, acetate, butyrate, hydrogen gas and CO_2_) with ordinary differential equations assuming Monod kinetics (see Material and Methods). The model differentiated between substrates required for growth and co-substrates such as acetate that enhanced growth but were not required. It also took species-specific differences in lag phases into account. Since we observed that FP did not deplete fructose, presumably because of a lack of co-factors, we introduced a dependency on an undefined metabolite referred to as “unknown compound”.

We parameterized this model on selected mono-culture experiments and then predicted mono-culture dynamics (Figure 4 A-C, Supplementary Figure 1). The model reached high prediction accuracy for FP and RI, but did not describe well the experimental data for BH (see Table 1). More precisely, the model showed that BH did not consume formate and fructose as quickly as expected if its growth would follow Monod kinetics. We confirmed culture homogeneity by analyzing 16S rRNA gene sequencing data of the last sample (Supplementary Figure 2). A yeast contaminant (S. *cerevisiae* S288c) detected in RNA-seq data of BH mono-culture samples (Supplementary Figure 3) does not explain the incongruence between growth and energy source consumption, since *i)* no contamination was observed on plates inoculated with bioreactor samples and incubated in anaerobic and aerobic conditions, *ii) S. cerevisiae* would consume fructose and *iii)* no ethanol production was measured. We also found only small concentrations of potential peptide degradation products (isobutyric acid and isovaleric acid). We therefore assume that BH in mono-culture initially grew on undefined medium components and only later switched to formate and fructose, but the time resolution was too low to model this potentially biphasic growth. We also compared model performance for RI with and without product inhibition by hydrogen gas. Since we found no differences in model performance, we removed an initial hydrogen gas inhibition term.

**Figure 4:**
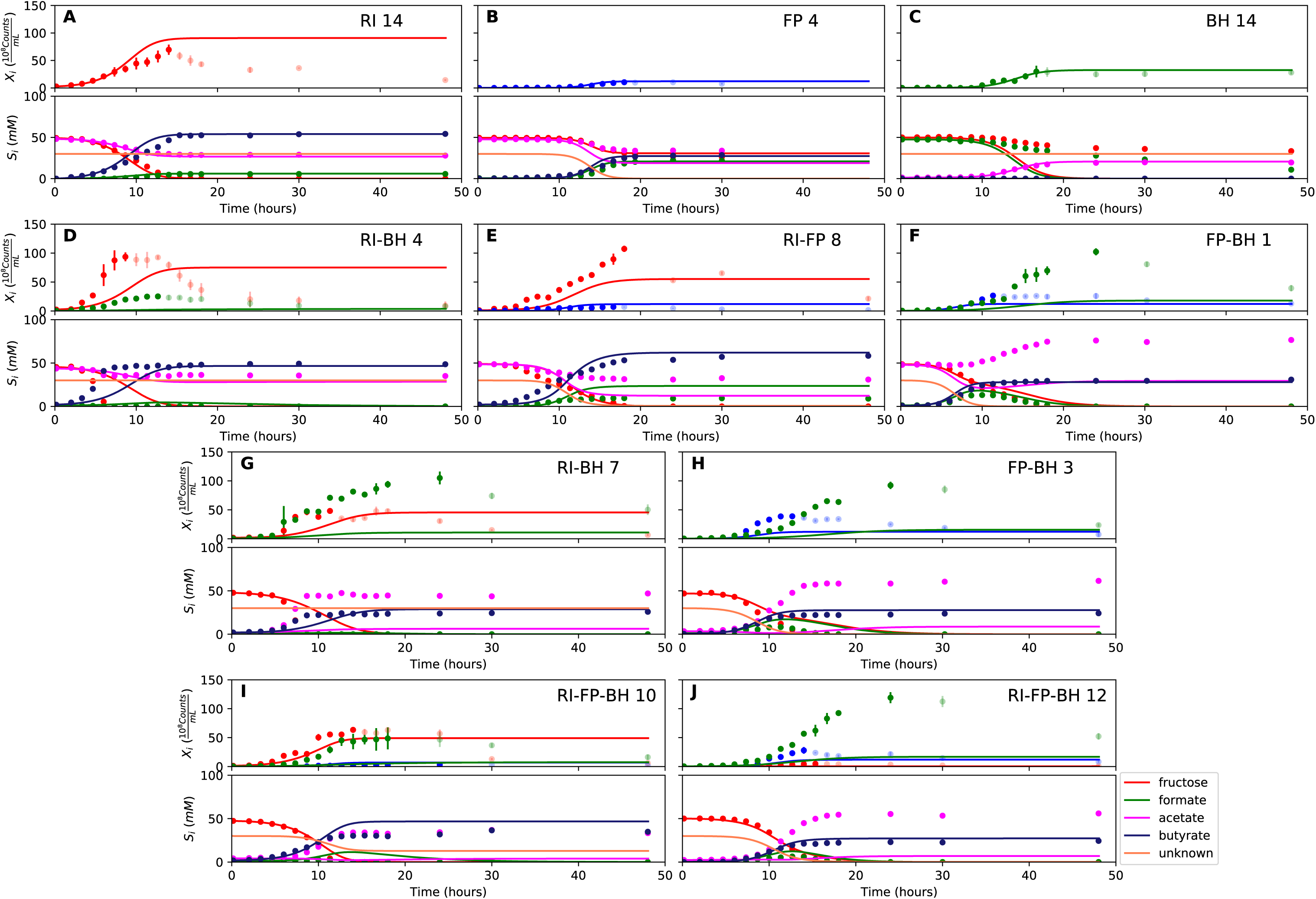
Model parameterized on mono-cultures does not fit co-culture data well. (A-C) Fit to mono-culture experiments selected for parameterization. (D-F) Fit to selected co-culture experiments with initial acetate. (G-H) Fit to selected co-culture experiments without initial acetate. (I-J) Fit to tri-cultures dominated by RI and BH versus FP and BH, respectively. Lines represent model predictions and dots represent observations. The whiskers represent technical variation across triplicates. The whiskers represent technical variation across triplicates. Transparent points indicate declining cell numbers; corresponding samples were not taken into account for model fitting. The unknown compound represents an unspecified co-substrate assumed to be required by *Faecalibacterium prausnitzii.* Metabolites not included in the model are omitted from the plot. Experiment identifiers indicate which of the biological replicates is displayed. The model was parameterized on experiments RI_8, RI_14, FP_4, FP_15 and BH_14.

### *Blautia* consumes formate produced by *Roseburia* and *Faecalibacterium*

When growing FP and BH together, we observed that fructose was entirely depleted and acetate, butyrate, hydrogen gas, CO_2_ and small concentrations of lactate were produced (Figure 3F). Interestingly, there was an initial production of formate, which was then consumed, confirming that formate was cross-fed from FP to BH. Formate consumption was also observed without initial acetate (Figure 3H).

In the bi-culture of RI and BH, CO_2_, hydrogen, butyrate and small concentrations of lactate were produced, whereas fructose and a small amount of acetate were consumed (Figure 3D). The same fermentation products were also obtained in the absence of initial acetate (Figure 3G). In contrast to RI in mono-culture, no formate was detected, suggesting that it was entirely cross-fed to BH. It was unclear whether CO_2_ and hydrogen gas produced by RI reached sufficiently high concentrations to be cross-fed to BH.

Finally, when RI and FP were co-cultivated, fructose and acetate were consumed and butyrate, formate, hydrogen gas and CO_2_ were produced (Figure 3E). The finding that formate reached lower concentrations than in FP mono-culture already hints at a negative effect of RI on FP.

### Comparison of mono- and co-culture data suggests ecological interactions

Since Gause’s early work on competition between yeast and *Paramecium* species (Gause, 1932, Gause, 1934), growth rates in mono- and bi-culture experiments have been compared to determine ecological interactions (e.g. (de Vos, Zagorski et al., 2017, Freilich, Zarecki et al., 2011, Wang, Wei et al., 2017)). The rationale is that growth rates in bi-culture should increase for mutualistic organisms as compared to mono-culture growth rates, whereas bi-culture growth rates should decrease for competitors.

When comparing maximal abundances, cross-feeding and competitive interactions were already apparent. Both FP and BH reached significantly higher maximal bacterial counts in FP-BH bi-cultures and in tri-cultures with FP dominance (Figure 3F, 3H and 3J) than they did in mono-culture (Figure 3B and 3C), suggesting a mutualistic relationship (unpaired two-sided Wilcoxon FP shift: 0.4, FP 95% confidence interval: 0.12 and 0.55, FP p-value: 0.03; BH shift: 0.5, BH 95% confidence interval: 0.33 and 0.69, BH p-value: 0.017). The maximal cell number of FP tended to be lower when competing with RI (Figure 3E) than when grown alone (unpaired two-sided Wilcoxon shift: 0.47, 95% confidence interval: −0.03 and 1.42, p-value: 0.11). Interestingly, there was no difference in maximal bacterial counts for RI alone versus grown with FP in bi-cultures and tri-cultures with RI dominance (unpaired two-sided Wilcoxon shift: 0.07, 95% confidence interval: −0.39 and 0.31, p-value: 0.69), so that formally, their relationship could be described as amensalism (one organism is affected negatively whereas the other is not affected). Finally, according to maximal bacterial counts, BH benefited more from the presence of FP than from RI (unpaired two-sided Wilcoxon shift: 0.29, 95% confidence interval: 0.06 and 0.93, p-value: 0.008).

### Model needs bi-culture data to accurately predict tri-culture dynamics

When growing all three gut bacterial strains together, fructose was consumed and butyrate, acetate, CO_2_, hydrogen gas as well as lactate were produced. Formate was produced initially, but quickly consumed (Figure 3I and 3J).

We performed the tri-culture six times with varying species proportions in the inoculum and found that in all tri-cultures, BH was always co-dominant, together with either RI or FP. In two out of the six cases, RI won over FP as the co-dominant partner of BH, whereas in the four other cases, FP won. The result mattered for the final butyrate concentrations, which averaged 37.5 mM when RI won and 23.5 mM when FP won.

We attempted to describe tri-culture dynamics with the model parameterized on mono-cultures, but failed to obtain a good fit (see Table 1 and Supplementary Figures 4 and 5). After a series of tests, we concluded that incorporating co-culture data was necessary to describe tri-culture dynamics. We finally selected two FP mono-cultures and the RI-BH and FP-BH bi-cultures with initial acetate to parameterize our model. As a validation, we predicted the behavior of RI-BH and FP-BH bi-cultures in the absence of initial acetate, which resulted in a good fit (Figure 5G and 5H, Supplementary Figure 7). The model parameterized on mono- and bi-cultures fitted the tri-culture data better than the model parameterized on mono-cultures only (Table 1, Figure 4I and 4J, Figure 5I and 5J, Supplementary Figures 5 and 8).

**Figure 5:**
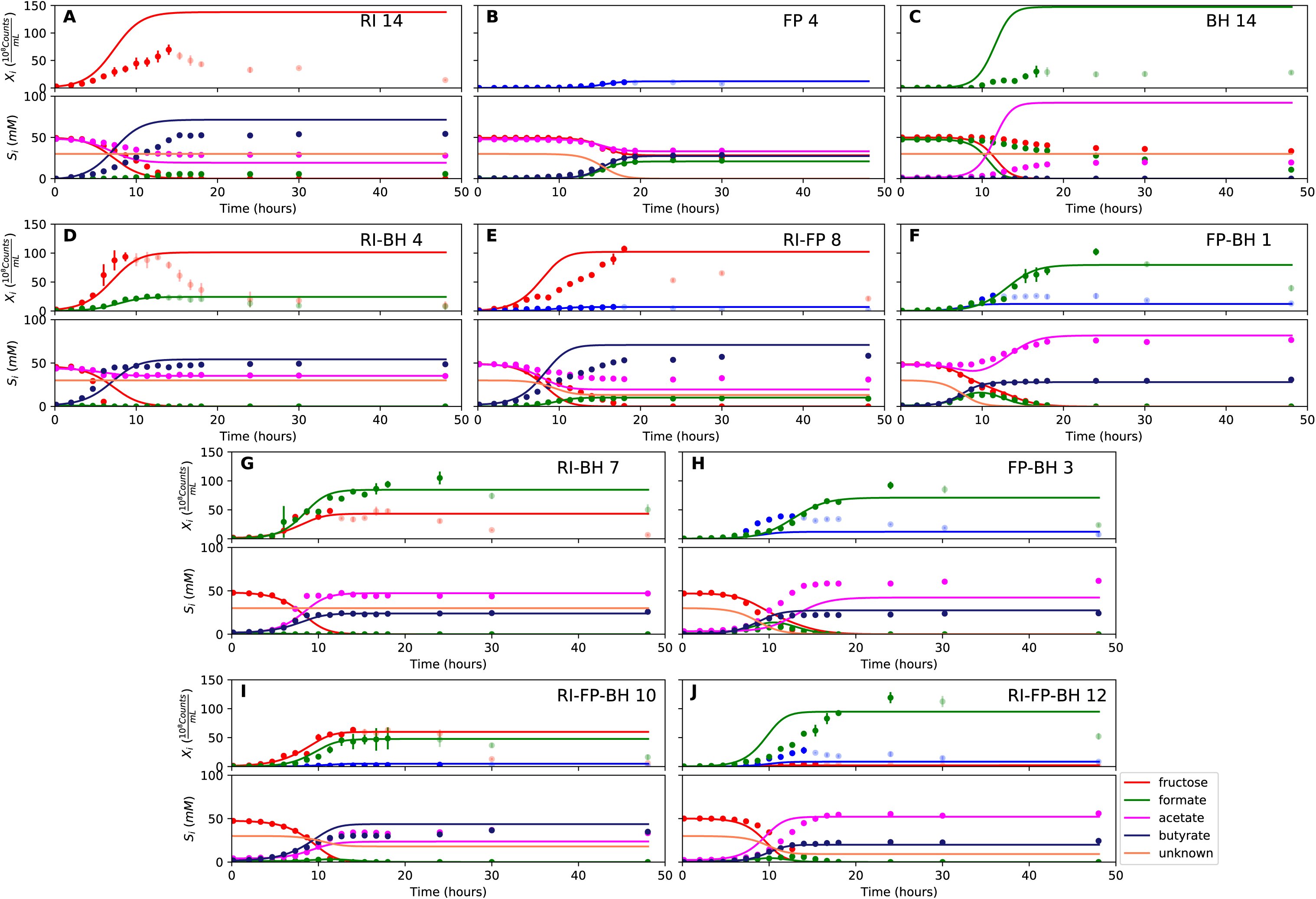
Model parameterized on mono- and bi-cultures improves fit to co-culture data as compared to parameterization on mono-cultures alone. (A-C) Fit to selected mono-culture experiments. (D-F) Fit to selected co-culture experiments with initial acetate (D and F were included in parameterization). (G-H) Fit to selected co-culture experiments without initial acetate, which were not part of the parameterization. (I-J) Fit to tri-cultures dominated by RI and BH versus FP and BH, respectively. Lines represent model predictions and dots represent observations. The whiskers represent technical variation across triplicates. The whiskers represent technical variation across triplicates. Transparent points indicate declining cell numbers; corresponding samples were not taken into account for model fitting. The unknown compound represents an unspecified co-substrate assumed to be required by *Faecalibacterium prausnitzii.* Metabolites not included in the model are omitted from the plot. Experiment identifiers indicate which of the biological replicates is displayed. The model was parameterized on experiments FP_4, FP_15, FP_BH_1, FP_BH_2 and RI_BH_4.

When inspecting the differences between the two parameterizations, we found that the model parameterized on mono-cultures predicted lower abundances for all three species in bi- and tri-cultures than they actually reached (Figure 4D-J, Supplementary Figures 4 and 5). Vice versa, the model parameterized on mono- and bi-cultures predicted too high abundances for RI and BH in mono-culture (Figure 5A and 5C, Supplementary Figure 6; the FP mono-culture was included in the parameterization). According to the higher maximal cell counts in tri-culture predicted with the second than the first parameterization (unpaired Wilcoxon: 0.002), BH did significantly better in tri-culture than expected based on its mono-culture growth.

The fact that the model describes well both mono- and tri-culture dynamics is a sign of emergent behavior in the presence of interaction partners. When looking at the parameters inferred from mono- and bi-cultures (given in Supplementary Table 2), BH’s consumption rates for formate and fructose and RI’s fructose consumption rate were lower compared to their values obtained from mono-culture parameterization, whereas their maximal growth rates were not much affected (BH) or increased (RI). Thus, according to this analysis, less of the energy source is needed in the presence of an interaction partner than in mono-culture.

### Initial abundance and lag phase predict species dominance in tri-culture

Next, we tested whether dominance in tri-culture could be predicted from lag phase and initial abundance. Towards this aim, we computed the FP/RI ratio in simulations with varying lag phase and initial abundance. Experimental observations of dominance agreed well with model predictions (Figure 6A and 6B). Our systematic investigation also showed that there was a non-linear relationship between initial RI abundance and FP dominance (Figure 6C-F). Thus, even when knowing initial abundances, lag phases and species interactions, it is hard to predict the winner (and thereby resulting butyrate concentration) intuitively without a model in hand. The final abundances of the three strains in simulations are also non-linearly dependent on other parameters, including BH’s growth rate, its fructose consumption rate and its fructose half-saturation constant (Supplementary Figure 9). These results underline that besides kinetic parameters, initial conditions and lag phase can determine strain abundances in co-culture in a non-linear way.

**Figure 6:**
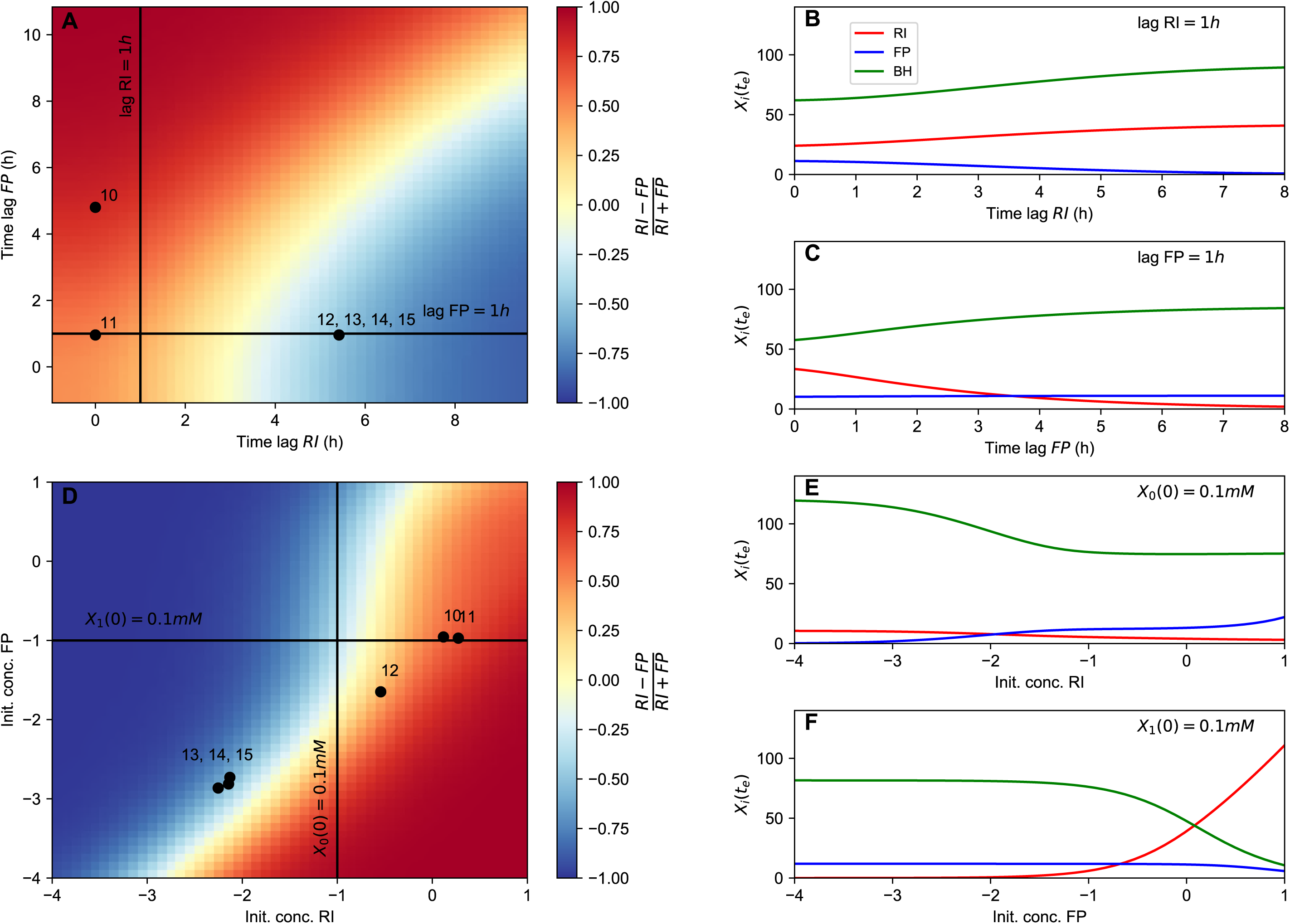
Initial conditions determine dominance in tri-culture. (A) The tri-culture dynamics is simulated with different lag phase values for *Faecalibacterium prausnitzii* (FP) and *Roseburia intestinalis* (RI) and the resulting end point abundance ratio of FP and RI is plotted in a heat map that is colored in blue for FP dominance and in red for RI dominance. The observed tri-culture data (black circles) are plotted according to the estimated experimental lag phases for RI and FP. The predicted RI or FP dominance agrees with the observed dominance in all six cases. (B) The tri-culture dynamics is simulated for varying initial abundances (init. conc.) of FP and RI and their resulting end point abundance ratio is visualized in a heat map. Three of the four FP-dominated experiments (13-15) and both RI-dominated experiments (10 and 11) are situated within their predicted region of dominance. (D-F) The dependency of the end point abundances of the three species on the lag phase and initial abundance of FP and RI is shown with simulations. This dependency is non-linear, especially for initial abundances of FP and RI, illustrating that dominance in batch is highly sensitive to initial conditions. The simulations were carried out with the model parameterized on mono- and bi-culture data. For the simulations in (A), the initial abundances of RI, FP and BH were set to 0.58, 0.04 and 0.21, respectively, whereas for the simulations in (B), the lag phases for RI, FP and BH were set to 0.33, 0.08 and 0.1, respectively. These initial abundance and lag phase values represent the averages of observed initial abundances and estimated lag phases across all tri-culture experiments.

### Altered gene expression in response to interaction partners provides first insights into emergent behavior

To further investigate the emergent behavior, we sequenced RNA for the three mono-cultures and the tri-culture where FP co-dominated for three time points and two biological replicates and assessed significantly differential gene expression across all samples in mono-versus tri-cultures for all three species (Supplementary Table 3). In total, 6.7%, 7.9% and 1.6% of RI’s, FP’s and BH’s proteins respectively were significantly differentially expressed (protein numbers taken from UniProt (The UniProt Consortium, 2017)). Interestingly, in tri-culture, FP down-regulated a series of enzymes needed for vitamin B_12_ coenzyme biosynthesis. Since cobalamin (vitamin B_12_) was one of the co-factors suspected to limit FP growth in mono-culture, this finding may mean that FP benefited from greater co-factor availability in tri-culture. In tri-culture, FP also up-regulated enzymes involved in amino acid and oligopeptide transport and amino acid and protein biosynthesis. BH likewise up-regulated amino acid biosynthesis in tri-culture. For RI, which reached lower abundances in the selected tri-cultures than in mono-culture, the transcription response was mixed: some amino acid biosynthesis enzymes were down-regulated, others up-regulated (including enzymes involved in ornithine biosynthesis). However, expression of ribosomal proteins was lower than in RI mono-culture, in agreement with its long lag phase in the selected tri-cultures. In summary, the analysis of differential gene expression uncovered a number of metabolic changes in the presence of interaction partners, thus supporting further the altered behavior detected through modeling.

## Discussion

Here, we investigated the dynamics of a well-defined small but representative gut microbial community *in vitro.* We found that BH was metabolically versatile and grew as fast as primary fermenters such as RI. We demonstrated experimentally that formate was cross-fed between BH on the one hand and FP and RI on the other and confirmed mutualistic as well as competitive interactions between these three bacterial strains. When growing on formate, we identified BH to be a net producer of both hydrogen and carbon dioxide, in contrast with its traditionally assumed role in the gut ecosystem. While formate is rarely highlighted as a key intermediate in gut cross-feeding interactions, it has been reported to be an end-product of primary polysaccharide degradation by both *Bifidobacterium* and *Lactobacillus spp.* (Falony, Lazidou et al., 2009b, Moens et al., 2017). Hence, our results invite a re-evaluation of the ecological niche of BH in relation with microbial formate production potential.

The model, which encodes our knowledge of the system, is not only important for predictions, but also as a reference. We gained insights from its agreements as well as from its disagreements with our observations. For instance, we assumed initially that RI would be inhibited by the hydrogen gas it generated. However, a hydrogen gas inhibition term was not required to accurately describe RI behavior in mono-culture, which implied that hydrogen gas inhibition did not affect RI growth at the concentrations reached in our experiments. We also needed the model to ascertain that changes from mono- to bi- or tri-cultures were not just due to variations in the inoculum composition or the lag phase, but that there was a true change in the dynamics that the model parameterized on mono-cultures alone could not capture. We confirmed this emergent behavior with RNA-seq, which revealed significantly different gene expression in tri-culture as compared to mono-culture, especially for FP and BH. The down-regulation of FP’s vitamin B12 coenzyme biosynthesis pathway in tri-culture is of particular interest, since it suggests that dependency on co-factors changes with interaction partners. It has been posited that the majority of gut microorganisms in need of B12 precursors is unable to synthesize them (Degnan, Taga et al., 2014). If this need is altered by the presence of interaction partners, it cannot be exploited as easily for selective manipulation as suggested in (Degnan et al., 2014).

Although species-level kinetic models parameterized on mono-cultures may in some cases describe bi-culture dynamics correctly (Van Wey, Cookson et al., 2014), our example shows that this is not a general property. This means that models of multi-species communities will probably have to take the internal metabolism and its response to interaction partners into account. Gut bacteria such as BH have flexible metabolic strategies that they employ according to circumstances. Emergent metabolism in the presence of interaction partners has been described in theoretical work before (Chiu, Levy et al., 2014), but has been rarely investigated experimentally ((Aharonovich & Sher, 2016)). Constraint-based modeling approaches, which take the entire metabolism into account (Orth, Thiele et al., 2010), require high-quality metabolic reconstructions for each community member, which take months of curation effort to obtain (Thiele & Palsson, 2010). Thus, scaling species-level quantitative models to larger communities will be a formidable challenge.

Mono- and bi-cultures are increasingly carried out in batch in a high-throughput fashion to determine ecological interactions and to quantify their strengths (de Vos et al., 2017, Sher, Thompson et al., 2011). Such systematic quantification is an important step forward, but there are challenges to tackle. Our work showed that dominance in batch may sensitively (i.e. non-linearly) depend on initial conditions such as the lag phase and the initial abundance, both of which are hard to control experimentally. Thus, a growth experiment performed in biological replicates but with the same inoculum may identify one species as the winner and another as the loser. However, a replicate with a slightly different inoculum composition may reach the opposite conclusion. Such a dependency on the initial conditions (albeit with larger abundance differences) has also been reported in several competition experiments for *Streptomyces* species (Wright & Vetsigian, 2016) and may thus be a common case. To ascertain that bacteria change their behavior in response to an interaction partner, RNA-seq can be carried out on mono- and bi-cultures (Aharonovich & Sher, 2016, Plichta, Juncker et al., 2016). Here, we showed that a model can also reveal emergent behavior by its failure to describe co-culture dynamics when parameterized on mono-cultures only. It is an open question how to scale up such approaches to achieve the high-throughput needed for systematic measurements of interaction strengths.

This is to the best of our knowledge the first *in vitro* investigation of a defined bacterial community that combines mutualism with competition in two cases. In the case of FP and BH, mutualism appears to supersede competition, leading to increased maximal bacterial numbers coupled with up-regulation of biosynthesis for both interaction partners. For RI and BH, we had no such clear experimental evidence, since RNA-seq was performed on tri-cultures dominated by FP, but the comparison of maximum bacterial numbers across mono-, bi- and tri-cultures suggests that RI and BH do not benefit as much from each other as FP and BH do. Since the model described RI-BH bi-culture dynamics well without taking CO_2_ and hydrogen gas cross-feeding into account, we assume that due to their low partial pressure gasses are less efficiently cross-fed to BH than formate, although both are likely metabolized via the same pathway (Wood-Ljungdahl). Thus, interactions that look similar on paper can play out differently, depending on the environment.

In the two replicates of the RI-FP bi-culture, RI and FP both survived (albeit FP in far lower numbers) despite competing for the same substrate, presumably because in our experimental set-up, the time until nutrient depletion was too short for the competitive exclusion principle to apply.

Our tri-culture experiments also demonstrated the importance of initial conditions on fermentation end-products. According to our model, the initial abundance and lag phase determined whether butyrate reached high or low concentrations in the tri-culture fermentations. Since these are likely to be relevant parameters in the gut environment and difficult to control, cocktail communities will have to be designed such that they will carry out their function across a wide range of initial conditions.

While our work highlighted a number of challenges to microbial community modeling, the model’s ability to predict tri-culture dynamics from bi-cultures gives hope that with sufficient knowledge, we will ultimately be able to model more complex microbial communities.

## Material and Methods

### Microorganisms and media

Human isolates of *Roseburia intestinalis* A2-165 (DSM 14610^T^), *Faecalibacterium prausnitzii* L1-82 (DSM 17677^T^) and *Blautia hydrogenotrophica* S5a33 (DSM 10507^T^) (abbreviated RI, FP and BH, respectively) were obtained from the Deutsche Sammlung von Mikroorganismen und Zellkulturen (DSMZ, Germany) and stored at −80 °C in reinforced clostridial medium (RCM; Oxoid Ltd., Basingstoke, United Kingdom), supplemented with 25% (vol/vol) of glycerol as a cryoprotectant.

A recently published medium for colon bacteria (mMCB) that allows growth of FP (Moens et al., 2016) was modified by adding nitrogen sources and trace elements as detailed below. This medium had the following composition (concentrations in g L^−1^): bacteriological peptone (Oxoid), 6.5; soy peptone (Oxoid), 5.0; yeast extract (VWR International, Darmstadt, Germany), 3.0; tryptone (Oxoid), 2.5; NaCl (VWR International), 1.5; K_2_HPO_4_ (Merck, Darmstadt, Germany), 1.0; KH_2_PO_4_ (Merck), 1.0; Na_2_SO_4_ (VWR International), 2.0; MgSO_4_⋅7H_2_O (Merck), 1.0; CaCl_2_⋅2H_2_O (Merck), 0.1; NH_4_Cl (Merck), 1.0; cysteine-HCl (Merck), 0.4; NaHCO_3_ (VWR International), 0.2; MnSO4^⋅^H_2_O (VWR International), 0.05; FeSO_4_·7H_2_O (Merck), 0.005; ZnSO_4_⋅7H_2_O (VWR International), 0.005; hemin (Sigma-Aldrich, Steinheim, Germany), 0.005; menadione (Sigma-Aldrich), 0.005; and resazurin (Sigma-Aldrich), 0.001. The medium was supplemented with 1 mL L^−1^ of selenite and tungstate solution (NaOH (Merck), 0.5; Na_2_SeO_3_·5H_2_O (Merck), 0.003; Na_2_WO_4_⋅2H_2_O (Merck), 0.004 and 1 L distilled water) and 1 mL L^−1^ of trace element solution SL-10 [HCl (Merck, 25%, vol/vol; 7.7 M), FeCl_2_⋅4H_2_0 (Merck), 1.5; ZnSO_4_⋅7H_2_O (VWR International), 0.148; MnSO4⋅H_2_O (VWR International), 0.085; H_3_BO_3_ (Merck), 0.006; CoCl_2_·6H_2_0 (Merck), 0.19; CuSO_4_⋅5H_2_0 (VWR International), 0.0034, NiCl_2_⋅6H_2_0 (Merck), 0.024, and Na_2_MoO_4_⋅2H_2_0 (Merck), 0.036). Acetate (50 mM or 6.8 g L^−1^ of CH_3_COO^−^Na^+^3H_2_O; Merck) was added to the medium for the monoculture fermentations with RI and FP, whereas formate (50 mM or 3.4 g L^−1^ of HCOO^−^Na^+^; VWR International) was added to the medium for the monoculture fermentations with BH. The pH of the medium was adjusted to 6.8 and the medium was autoclaved at 210 kPa and 121 °C for 20 min. After sterilization, D-fructose (Merck) was added as the sole energy source aseptically, at a final concentration of 50 mM fructose using sterile stock solutions obtained through membrane filtration using Minisart filters (pore size, 0.2 μm; Sartorius, Göttingen, Germany).

### Cultivation experiments in stationary bottles

Monoculture cultivation experiments for BH were performed in stationary glass bottles without controlling the pH (screening). The bottles contained 50 mL of heat-sterilized pH 6.8 mMCB medium, supplemented with either 50 mM of D-fructose (Merck), D-glucose (Merck), D-galactose (Merck), L-fucose (Merck), sodium formate (VWR International), sodium acetate trihydrate (Merck), DL lactic acid (VWR International), oligofructose (Raftilose P95; Beneo-Orafti NV, Tienen, Belgium; (Falony et al., 2009b)) or inulin (OraftiHP; Beneo-Orafti; (Falony et al., 2009b)) as the sole energy sources. Additional cultivation experiments were performed in medium devoid of any main energy source to test autotrophic growth. For the cultivation experiments in bottles, stock solutions of fructose, glucose, galactose, fucose, sodium formate, sodium acetate trihydrate, and lactic acid were initially made anaerobically through autoclaving at 210 kPa and 121 °C for 20 min. The solutions were subsequently filter-sterilized and transferred into glass bottles, which were sealed with butyl rubber septa that were pierced with a Sterican needle (VWR International) connected with a Millex-GP filter (Merck) to assure sterile conditions. For the cultivation experiments with lactate, the pH was adjusted to 6.8 under anaerobic conditions, using sterile solutions of sodium hydroxide (Merck). Stock solutions of oligofructose and inulin were made sterile by membrane filtration. The inocula were prepared as follows. Cells of the strains under study were transferred from −80°C to test tubes containing 10 mL of RCM that were incubated anaerobically at 37 °C for 24 h. Subsequently, the strains were propagated for 12 h in glass bottles containing 50 mL of heat-sterilized pH 6.8 mMCB medium, supplemented with the energy source under study, always at a final concentration of 50 mM fructose equivalents. These pre-cultures were finally added to the glass bottles aseptically. During the inoculum build-up, the transferred volume was always 5% (vol/vol). All bottles were incubated anaerobically at 37°C in a modular atmosphere-controlled system (MG anaerobic work station; DonWithley Scientific Ltd., West Yorkshire, United Kingdom) that was continuously sparged with a mixture of 80% N_2_, 10% CO_2_, and 10% H_2_ (Air Liquide, Paris, France). Samples for further analyses were withdrawn after 0, 6, 12, 24, 48, and 100 h. All experiments were performed at least in duplicate.

### Fermentation experiments

To prepare inocula, cells were transferred from −80 °C to test tubes containing 10 mL of RCM, and incubated at 37 °C for 24 h. Subsequently, the strains were propagated twice for 12 h in glass bottles containing 100 mL of mMCB medium (with acetate in the case of RI and FP, and with formate in the case of BH), supplemented with fructose. All incubations were performed anaerobically in a modular atmosphere-controlled system (MG anaerobic workstation) that was continuously sparged with a mixture of 80% N_2_, 10% CO_2_, and 10% H_2_ (Air Liquide). The inocula were finally added aseptically to the fermentors. During the inoculum build-up, the transferred volume was always 5% (vol/vol). Fermentations were carried out in 2-L Biostat B-DCU fermentors (Sartorius) containing 1.5 L of mMCB medium supplemented with the co-substrates (acetate and/or formate) if necessary and 50 mM of D-fructose as the energy source. Anaerobic conditions during fermentations were assured by continuously sparging the medium with N_2_ (PraxAir, Schoten, Belgium) at a flow rate of 70 mL min^−1^. The fermentation temperature was kept constant at 37 °C. A constant pH of 6.8 was imposed and controlled automatically, using 1.5 M solutions of NaOH and H_3_PO_4_. To keep the medium homogeneous, a gentle stirring of 200 rpm was applied. Temperature, pH, and agitation speed were controlled online (MFCS/win 2.1 software, Sartorius). Fermentations were followed for 48 h, with samples taken at 10 min and 2 h, 3 h, 5 h, 6 h, 7 h, 9 h, 10 h, 11 h, 13 h, 14 h, 15 h, 17 h, 18 h, 24 h, 30 h and 48 h after inoculation. At selected time points (3 h, 9 h and 15 h after inoculation), subsamples were treated for RNA extraction by adding 5 vol of RNAlater® (xxx company).

All mono and tri-culture fermentations were performed in triplicate. All bi-culture fermentations were performed in duplicate, except for the bi-culture fermentations using medium lacking acetate with FP and BH that were performed only once.

### Quantification of bacterial abundance

During all experiments, the optical density at 600 nm (OD_600_) was measured against ultrapure water as blank with a VIS spectrophotometer (Genesys 20; Thermo Scientific, Waltham, MA, USA). Each measurement was performed in triplicate. Total bacterial abundance was also measured by flow cytometry, using an Accuri C6 flow cytometer (BD Biosciences, Erembodegem, Belgium), as described previously (Moens et al., 2016). All samples were diluted in filter-sterilized water (Vittel, France) to obtain a concentration between 1.0 × 10^3^ and 5.0 × 10^6^ cells mL^−1^. Flow cytometric analysis was performed by mixing 500 μL of sample with 5 μ.L of a 100× SYBR Green I solution (Sigma-Aldrich) and 5 μL of a 500 mM ethylenediaminetetra acetic acid (EDTA) solution (Sigma-Aldrich). Afterwards, samples were left in the dark at room temperature for 15 min. Flow cytometric counts were obtained using an Accuri C6 flow cytometer (BD Biosciences), equipped with a 50 mW solid state laser (488 nm). Green fluorescence was measured in the FL1 channel (530 ± 15 nm) and all data were processed with the Cflow Plus software (Accuri). Gating was performed to distinguish signals from noise. All data were collected as a FL1/SSC density plot with a primary threshold of 10,000 on the FL1 channel. Measurements were performed in triplicate.

qPCR assays with species-specific TaqMan primers and probes were performed to quantify the abundance of each species separately. For this, 2 mL of fermentation sample was centrifuged at 20,570 × *g* for 20 min. Cells were washed in 2 mL of physiological solution (NaCl, 8.5 g L^−1^) and centrifuged again at 20,570 × *g* for 20 min to obtain washed cell pellets. Subsequently, these cell pellets were resuspended in 2 mL of physiological solution and diluted 20 times for DNA extraction. Direct DNA extractions by alkaline thermal lysis were performed based on (Girish, Haunshi et al., 2013, Rudbeck & Dissing, 1998), modified as follows: 100 μL of the sample was mixed with 100 μL of 0.2 M NaOH in a sterile micro centrifuge tube. The mixture was vortexed and heated at 90°C for 10 min, after which eight volumes (1600 μL) of 0.04 M Tris HCl pH 7.5 (Thermo Fisher Scientific) was added for pH neutralization. 4 μL of final mixture was used for qPCR. The extracted genomic DNA was stored at −20°C until qPCR amplification.

Calibration curves were obtained by initially growing all strains in RCM for 24 h, and two-fold propagation in medium for 12 h, as described above. From each of these grown cultures, separate fourfold decimal and nine-fold binary dilution series were prepared. The generation of cell pellets, direct extraction of DNA, and subsequent quantification of cell concentrations by flow cytometry were performed as described above, with the exception that, prior to DNA extraction, samples for calibration were diluted less than fermentation samples, to accommodate a wider qPCR quantification range.

Primers and oligoprobes (Supplementary Table S4) were manually designed using the online Primer3Plus software (Untergasser, Nijveen et al., 2007) and the genome sequences of the strains. Melting temperatures and presence of hairpins, self-dimers, and pair-dimers were double-checked using the online OligoCalc software (Kibbe, 2007). Secondary structures of the generated amplicons was investigated using the online Mfold program (Zuker, 2003). Primers and probes were synthesized by Thermo Fisher Scientific. Strain specificity of primers and probes was confirmed *in silico* by Primer-BLAST (Ye, Coulouris et al., 2012) and *in vitro* by PCR and qPCR analysis on genomic DNA of the strains (Supplementary Table S1). qPCR assays were carried out using a 7500 FAST Real-Time PCR system (Applied Biosystems, Carlsbad, CA, USA), equipped with 96-well plates. Each qPCR assay mixture of 20 μL contained 10.0 μL of TaqMan® Fast Universal PCR Master Mix (2X), no AmpErase® UNG (Thermo Fisher Scientific), 2.0 μL of each primer (3.0 μM), 2.0 μL of the TaqMan probe (1.5 μM), and 4.0 μL of extracted genomic DNA solution or sterile nuclease-free water (Thermo Fisher Scientific). The qPCR amplification program consisted of an initial denaturation step at 95°C for 20 s, followed by 45 two-step cycles at 95°C for 3 s and at 60°C for 30 s. In each run, negative (sterile nuclease-free water without genomic DNA) and positive controls (with extracted genomic DNA from the relevant strains) were used. The cycle threshold (Ct) values were determined using the automatically determined thresholds from the 7500 software v2.0.6 (Applied Biosystems). Finally, during a re-analysis of all qPCR runs, Ct values were normalized using an inter-plate calibrator to account for differences among qPCR runs. The above-described generation of cell pellets, direct extraction of DNA, and qPCR assays were performed in triplicate.

Contamination was checked by aerobic and anaerobic plating on RCM agar and 16S rRNA gene amplicon sequencing of end point fermentation samples (48 h). Sequencing was performed as described previously (D’hoe, Conterno et al., 2018).

### Metabolite profiling

Concentrations of fructose, as well as concentrations of formate, acetate, butyrate, lactate, and ethanol were determined through high-performance liquid chromatography (HPLC) with refractive index (RI) detection, using a Waters chromatograph (Waters, Milford, MA, USA) equipped with an ICSep ICE ORH-801 column (Transgenomic North America, Omaha, NE, USA), and applying external standards, as described previously (Falony et al., 2009b). Briefly, the mobile phase consisted of 5 mM H_2_SO_4_ at a flow rate of 0.4 mL min^−1^. The column temperature was kept constant at 35°C. Sample preparation involved a first centrifugation (4,618 × *g* for 20 min at 10°C) for removal of cells and debris, followed by the addition of an equal volume of 20% (mass/vol) trichloroacetic acid for protein removal. For determining oligofructose and inulin consumption, samples were incubated at room temperature for 24 h to assure complete hydrolysis of polysaccharides. Subsequently the samples were centrifuged (21,912 × *g,* 20 min, 4°C) and filtered (pore size of 0.2 μm; Uniflo 13 Filter Unit; GE Healthcare, Little Chalfont, United Kingdom), prior to injection (30 μL) into the column. Samples were analyzed in triplicate.

For an additional screening experiment with BH grown in the presence of 350 mM formate, the concentrations of ethanol, acetoin, acetic acid, propionic acid, butyric acid, isobutyric acid and isovaleric acid produced were determined via gas chromatography with flame ionization detection (GC-FID), using a FocusGC chromatograph (Interscience, Breda, The Netherlands) equipped with a Stabilwax-DA column (Restek, Bellefonte, PA, USA), and applying external standards, as described previously (Moens, Lefeber et al., 2014). The samples were analyzed in triplicate.

### Model definition

We modeled change of species abundances over time with the following three ordinary differential equations:

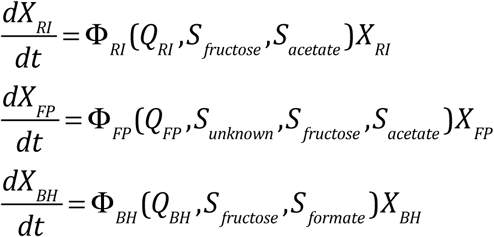

where *X* denotes species abundance, *S* metabolite concentration and *Q* a lag phase parameter.

The growth rates are then defined as nonlinear growth functions as in (Grivet, 2001, Smith & Waltman, 1995), which assume Monod kinetics (Monod, 1950):

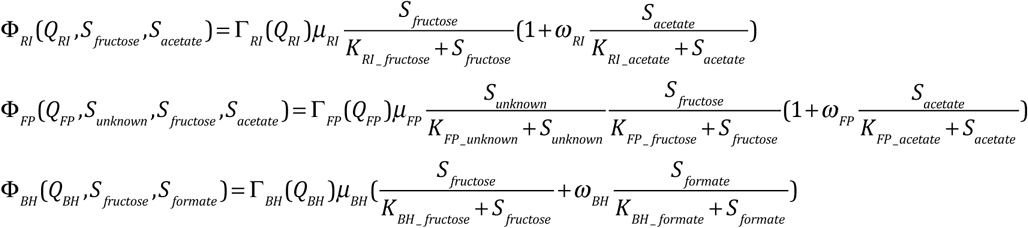

where *K* is the Monod (half-saturation) constant, μ is the maximal specific growth rate andω a weight parameter. Nutrient dependency can be either obligatory (growth without nutrient is not possible) or facultative (growth without nutrient is possible). For instance, the fructose uptake is multiplied with RI’s maximal growth rate, whereas its acetate uptake is modeled with an additive term. Therefore, in the absence of fructose, RI’s growth rate is zero, but not in the absence of acetate. The weight parameter adjusts how strongly a facultative substrate contributes to the overall growth rate. The unknown compound models the dependency of FP on an undetermined co-factor.

The lag phase function is defined as in (Baranyi & Roberts, 1994):

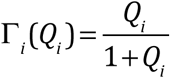 where *i* stands for RI, FP or BH.

The Q_i_ variables follow exponential growth:

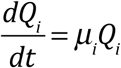

Thus, the larger the initial value of Q_i_, the shorter the lag phase.

The changes of metabolite concentrations are then modeled as follows:

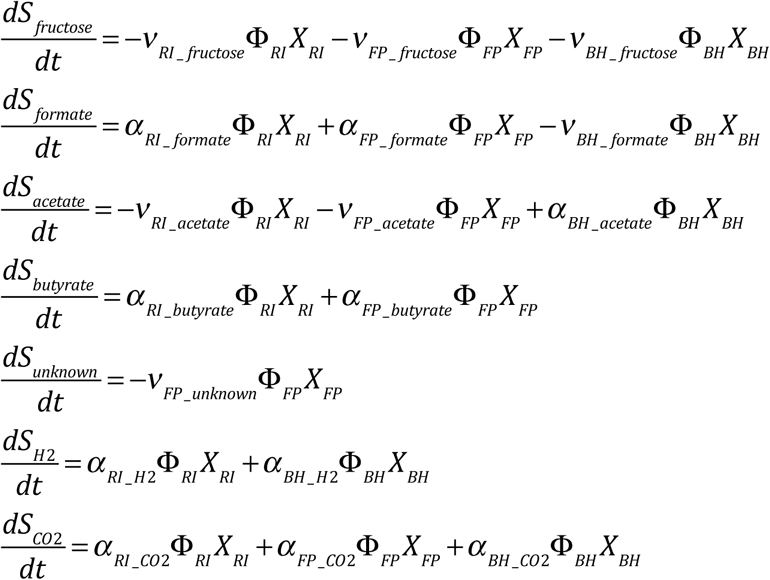

The *α* and *ν* parameters are production and consumption rates, respectively. Species abundance is measured in 10^8^ bacterial counts/mL, metabolite concentration in mM, the unit of *μ* is 1/h, the unit of K is mM, the unit of *α* and of *ν* is mM/(10^8^ bacterial counts/mL) and *ω* is dimensionless.

The model assumes that death rates are negligible. CO_2_ and hydrogen consumption by BH is not included in the final version of the model. We tried to account for CO_2_ consumption with a multiplicative term in BH’s growth rate. However, this did not improve the model fit. Since the model without CO_2_ describes RI-BH bi-culture dynamics well, we assume that BH grows mostly heterotrophically on fructose and on the formate produced by RI and that the hydrogen gas and CO_2_ produced by RI did not reach sufficient concentrations in the head space to allow autotrophic growth as observed during the screening, where the atmosphere contained 10% of hydrogen gas and 10% of CO_2_. The model definition is available as Source Code File in python (Model definition).

### Model parameterization

We parameterized our model on mono-cultures alone (parameterization 1) and on mono- and bi-cultures (parameterization 2). The goodness of fit of both parameterizations are summarized in Table 1.

The model was fitted using the function fmin() from the scipy Python package (Jones, Oliphant et al., 2001), to minimize the normalized root mean square error. An initial estimate of the parameters was obtained by manually fitting the data iteratively. The initial concentration of the unknown compound was set to 30 mM. Samples after the end of the log phase, when the bacterial counts started to decline, were omitted from the fitting. Parameterization 2 consisted of several steps, as fitting all parameters at once did not lead to convergence, due to the nonlinear growth rates. For this, FP was first fitted on two FP mono-cultures and cross-validated on the third one. The consumption parameters of BH were obtained from FP-BH bi-cultures with initial acetate, afterwards the maximal specific growth rates and half-saturation constants were obtained from the same bi-cultures. RI was fitted on a RI-BH bi-culture with acetate. Lag phases were calculated as the time to reach Γ_*i*_ (*Q*_*i*_) = 0.5:

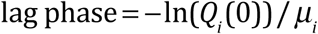

*Q*_*i*_*(0)* was estimated by visual inspection of the log plots. Model parameters obtained and maximal abundances predicted with both parameterizations as well as estimated lag phases are provided in Supplementary Table 2. Data and model fits were plotted with Python’s matplotlib (Hunter, 2007).

### RNA extraction and sequencing

Total RNA was extracted from RNAlater®-treated samples using the phenol-free total RNA purification kit coupled with DNase I treatment (VWR International) according to the manufacturer’s protocol for Gram-positive bacteria. A secondary DNAse digestion was performed using the Ambion® TURBO DNA-free™ DNase Treatment and Removal Reagents Kit (Thermo Fisher Scientific), after which the samples were purified using the RNA Clean & Concentrator™-25 kit (Zymo Research, Irvine, CA, USA) according to the manufacturer’s instructions.

The eluted RNA was stored at −80°C. The absence of DNA contamination was evaluated using PCR (35 or 40 cycles) and gel electrophoresis. The concentrations of the samples were determined with a Nanodrop, and with a Qubit 2.0 fluorometer using the Qubit dsDNA HS Assay Kit (Thermo Fisher Scientific). RNA integrity, expressed as the RNA integrity number (RIN), and yield were determined using RNA Nano/Pico 6000 LabChips (Agilent Technologies, Santa Clara, CA, USA) that were run in an Agilent 2100 Bioanalyzer (Agilent Technologies). Whereas most of the RINs were above 7, RINs of three BH mono-culture samples at 3 h, 9 h and 15 h were around 2.6 and in four cases (BH mono-culture at 15 h, FP mono-culture at 9 h, and for both tri-culture replicates at 3 h) the RINs could not be determined. However, by pooling over three extraction rounds, sufficient RNA for sequencing (minimum of 536 ng and median of 2800 ng) was obtained for all samples.

Library preparation encompassed the use of Ribozero rRNA removal for Gram-positive Bacteria and the Illumina TruSeq stranded mRNA Library preparation kit (IIlumina, San Diego, CA, USA). Library preparation was performed without the mRNA purification step, according to the manufacturer’s instructions. The enriched libraries were sequenced on an Illumina NextSeq 500 instrument (paired-end, 2×76 bp reads, Mid output kit, Illumina). From the Illumina platform, paired-end reads in FASTQ format (CASAVA 1.8, Phred + 33) were obtained and separated into distinct files for each single-end read and for each sample.

### RNA-seq analysis

The analysis of the raw sequencing reads was performed as follows: reads were trimmed using Trimmomatic (Bolger, Lohse et al., 2014) with the following parameters “CROP:74 HEADCROP:10 SLIDINGWINDOW:4:15 MINLEN:51”, to remove initial and last bases which had biases in their nucleotide content as reported by FastQC (Andrews, 2010), to remove stretches of low-quality bases and to keep reads with at least 51 bases after trimming. FastQC was re-run on the trimmed data to ensure the previous biases were corrected. SortMeRNA (Kopylova, Noé et al., 2012) was used with default parameters and included databases to remove rRNA reads.

With the remaining non-rRNA reads, we ran MetaPhlAn2 (Truong, Franzosa et al., 2015) with default parameters and database, and mash screen (Ondov & Philippy, 2017, Ondov, Treangen et al., 2016) with default parameters against the complete RefSeq genomes and plasmids database, to search for potential contaminants. Both MetaPhlAn2 and the top hits from mash screen found the correct bacterial genomes from the three strains used in this study, together with reads from yeast (*S*. *cerevisiae* S288c). Additionally, low amounts of the phage PhiX174 were reported by mash screen. To accurately quantify the presence of these potential contaminants in our samples, and to quantify gene expression from the cultured bacteria, we mapped the non-rRNA reads to these five species using Bowtie2 (Langmead & Salzberg, 2012) with default parameters except for “-X 800” to allow for longer insert sizes. The reference genomes used are the following: GCF_000156535.1_ASM15653v1_genomic.fna (RI), GCF_000157975.1_ASM15797v1_genomic.fna (BH), GCF_000162015.1_ASM16201v1_genomic.fna (FP), CF_000146045.2_R64_genomic.fna (yeast) and NC_001422.1 (PhiX174). Gene expression was quantified using the htseq-count Python script (Anders, Pyl et al., 2015) (with parameter −a 2 to exclude multimapping reads) for all species using their available *.gff reference annotation files. Given the small size of the PhiX174, we quantified the reads mapping to its entire genome rather than its gene expression.

Differential gene expression analysis of the three cultured strains was performed with DESeq2 (Love, Huber et al., 2014). To remove the effect of the different bacterial compositions in the tri-culture samples, we extracted the reads from each strain prior to the differential expression analysis, and analyzed each strain separately. In the DESeq2 design formula we included two factors: type of culture (mono- or tri-culture) and time (3 h, 6 h and 15 h). The results of the differential expression analyses were computed using a Wald test of the tri-culture versus the mono-culture samples.

For each strain, we extracted the genes significantly changing their expression (with Benjamini-Hochberg adjusted p-value < 0.05) in tri-culture and mapped them to different functional annotations downloaded from the IMG database (Markowitz, Chen et al., 2012): COG categories, COG numbers and KO numbers. The RNA-seq data processing code is available on GitHub (https://github.com/vllorens/syntheticGutCommunity).

## Acknowledgements

The technical assistance of ing. Wim Borremans for the metabolite analyses is gratefully acknowledged. We also thank Frédéric Leroy for helpful discussions.

## Availability of data and code

RNA-seq results have been deposited in the Short Read Archive under the study identifier SRP136465 (reviewer access: ftp://ftp-trace.ncbi.nlm.nih.gov/sra/review/SRP136465_20180410_112745_6a55596c15_df4993eea3b44eace1ee7f). Fermentation data have been submitted to Dryad (doi:10.5061/dryad.g83f29f, reviewer access: https://datadryad.org//review?doi=doi:10.5061/dryad.g83f29f). The model definition is provided in the Source Code File “Model definition”. The RNA-seq data processing code is provided on GitHub (https://github.com/vllorens/syntheticGutCommunity).

---

**Supplementary Figure 1.**
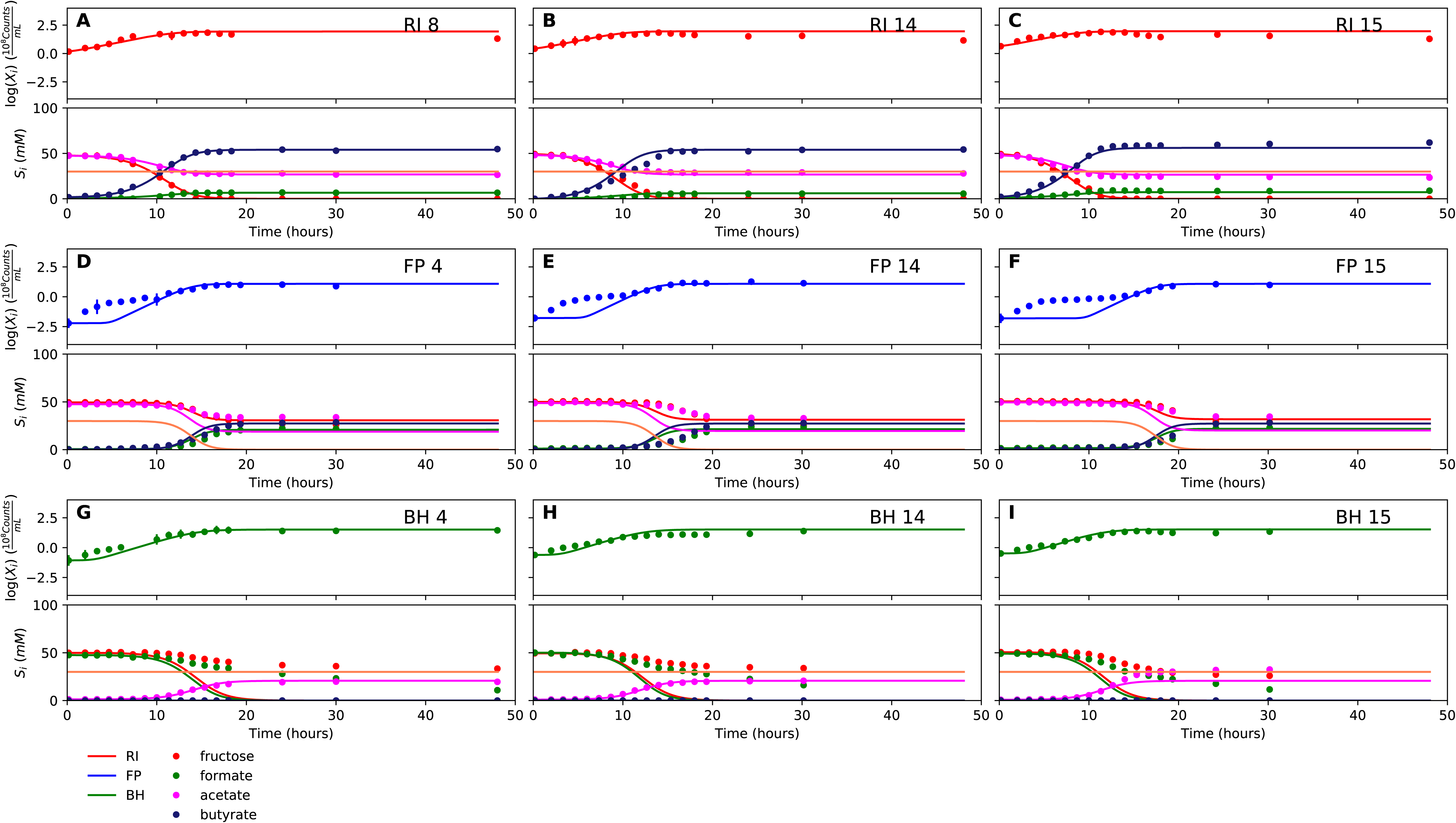
Fit to mono-culture experiments for the model parameterized on mono-cultures only. (A-C) Fit to *Roseburia intestinalis* mono-culture experiments. (D-F) Fit to *Faecalibacterium prausnitzii* mono-culture experiments. (G-I) Fit to *Blautia hydrogenotrophica* mono-culture experiments. Lines represent model predictions and dots represent observations. The whiskers represent technical variation across triplicates. The shaded regions indicate the length of the estimated species-specific lag phases. The unknown compound represents an unspecified co-substrate assumed to be required by *Faecalibacterium prausnitzii.* Metabolites not included in the model are omitted from the plot. Experiment identifiers indicate which of the biological replicates is displayed. The model was parameterized on experiments RI_8, RI_14, FP_4, FP_15 and BH_14.

**Supplementary Figure 2.**
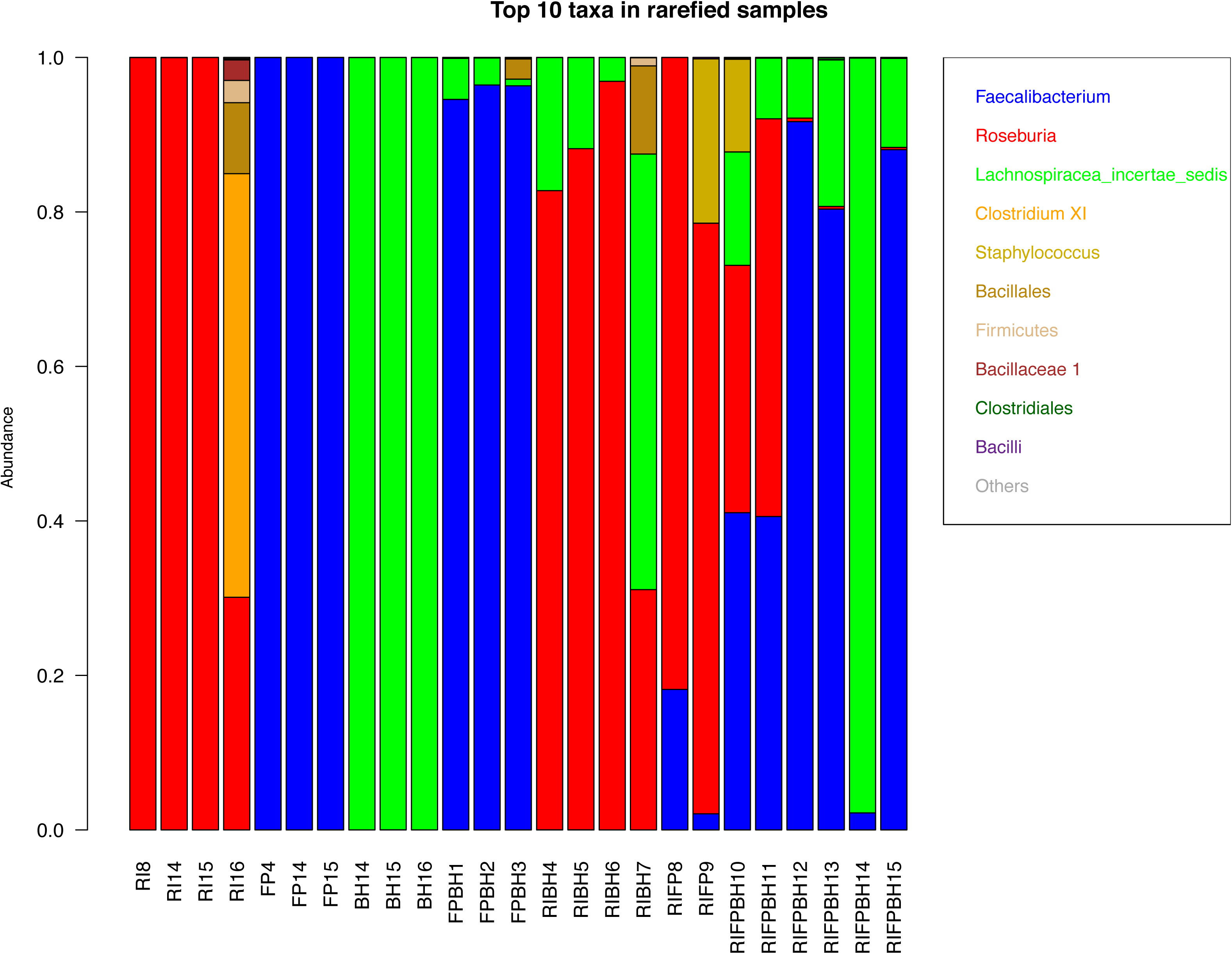
Test for prokaryotic contamination with 16S rRNA gene sequencing. For samples taken at the last time point, DNA was extracted and the V4 region of the 16S rRNA gene was amplified and sequenced. Raw reads are rarefied to 15,339 counts per sample and then converted into relative abundances. The top 10 taxa in each sample are shown. The abbreviations RI, FP and BH in sample identifiers stand for *Roseburia intestinalis, Faecalibacterium prausnitzii* and *Blautia hydrogenotrophica,* respectively. The taxon *Lachnospiraceae incertae sedis* contains BH. High relative abundances of potential contaminants (> 10%) were found in one RI monoculture (RI_16), one RI-FP co-culture (RI_FP_9) and one tri-culture (RI_FP_BH_10).

**Supplementary Figure 3.**
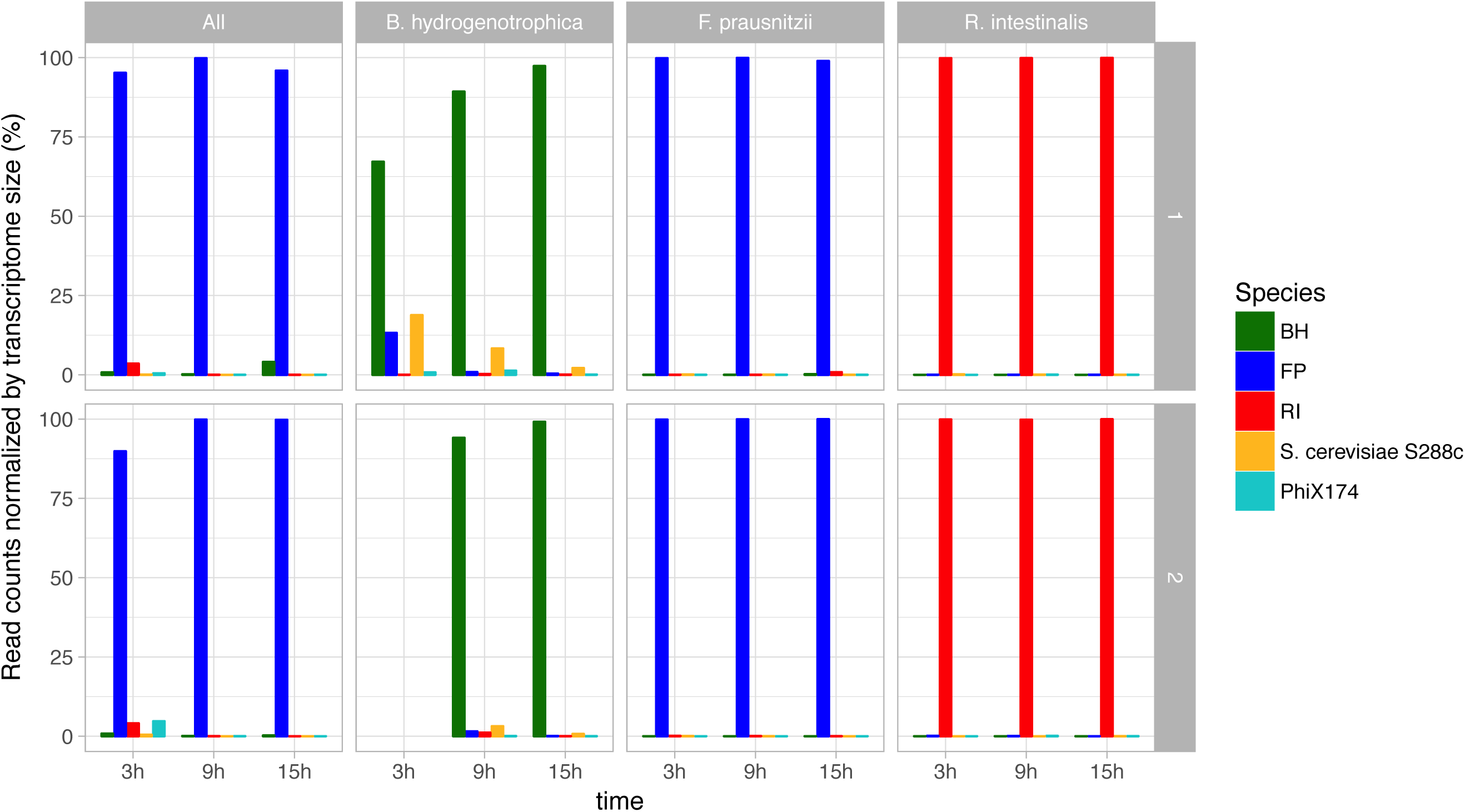
Test for viral, prokaryotic and eukaryotic contamination in RNA-seq data. The RNA of two non-prokaryotic organisms reached noticeable abundances: the bacteriophage phiX174, which is used as a control in Illumina sequencing, and the yeast *S. cerevisiae* S288c, which probably came from the yeast extract employed in the medium. In most of the samples, these potential contaminants have transcriptome-size-corrected abundances below 5%, but in one sample (*Blautia hydrogenotrophica* at 3 h) the yeast RNA abundance reached 18%. Taxonomic assignment was carried out with MetaPhlAn2 and mash screen against the complete RefSeq genomes and plasmids database. Total read counts were corrected for transcriptome size (genome size in the case of the bacteriophage). The bacterial contamination observed in the *Roseburia intestinalis* mono-culture with 16S rRNA gene sequencing was not confirmed with RNA-seq. Row 1 refers to experiments RI_16, FP_14, BH_16 and RI_FP_BH_14, while row 2 refers to experiments RI_15, FP_15, BH_15 and RI_FP_BH_15.

**Supplementary Figure 4.**
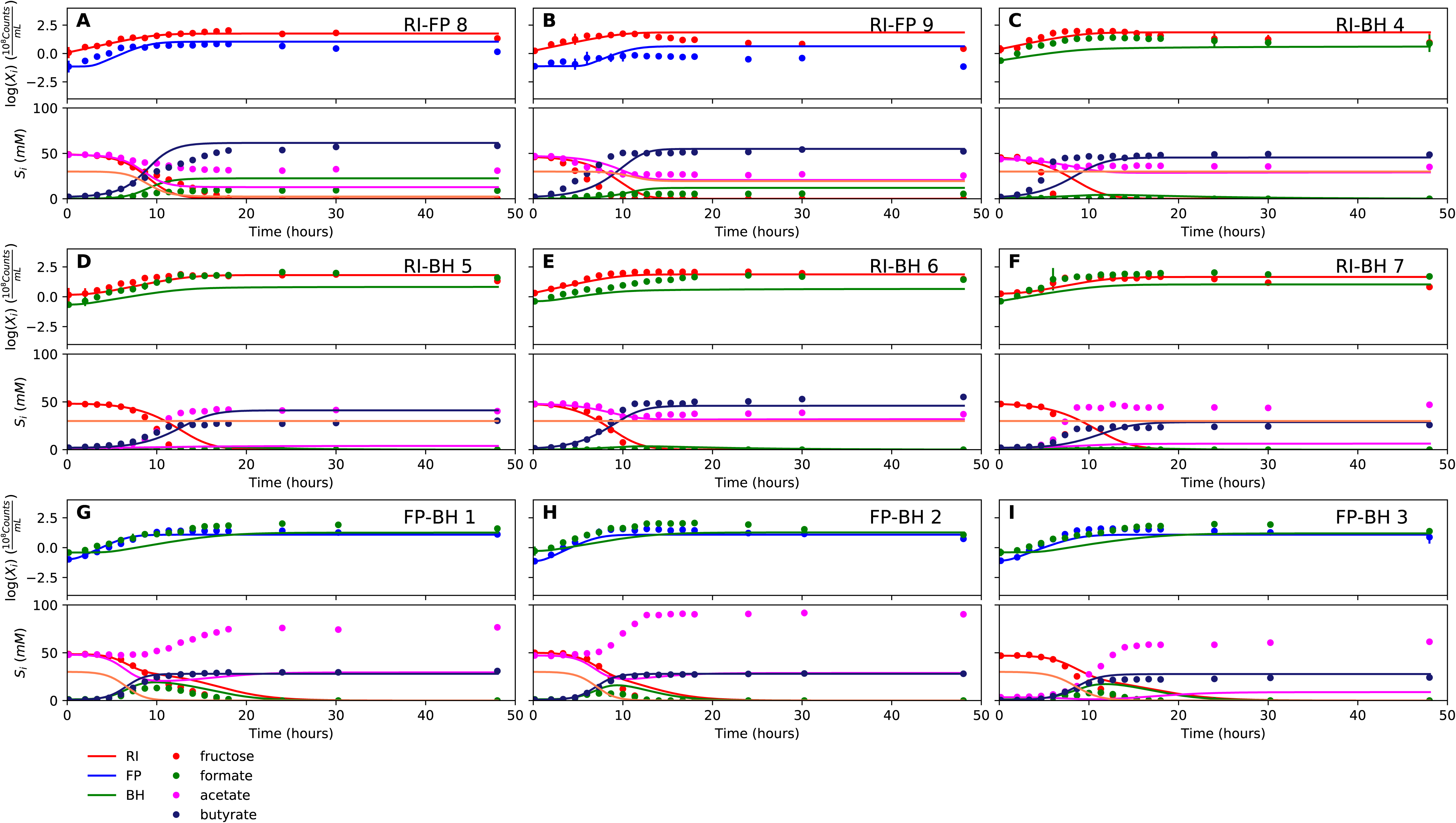
Fit to bi-culture experiments for the model parameterized on mono-cultures only. (A-B) Fit to *Roseburia intestinalis* and *Faecalibacterium prausnitzii* bi-culture experiments. (C-F) Fit to *Roseburia intestinalis* and *Blautia hydrogenotrophica* bi-culture experiments. (G-I) Fit to *Faecalibacterium prausnitzii* and *Blautia hydrogenotrophica* bi-culture experiments. Lines represent model predictions and dots represent observations. The whiskers represent technical variation across triplicates. The shaded regions indicate the length of the estimated species-specific lag phases. The unknown compound represents an unspecified co-substrate assumed to be required by *Faecalibacterium prausnitzii.* Metabolites not included in the model are omitted from the plot. Experiment identifiers indicate which of the biological replicates is displayed. The model was parameterized on experiments RI_8, RI_14, FP_4, FP_15 and BH_14.

**Supplementary Figure 5.**
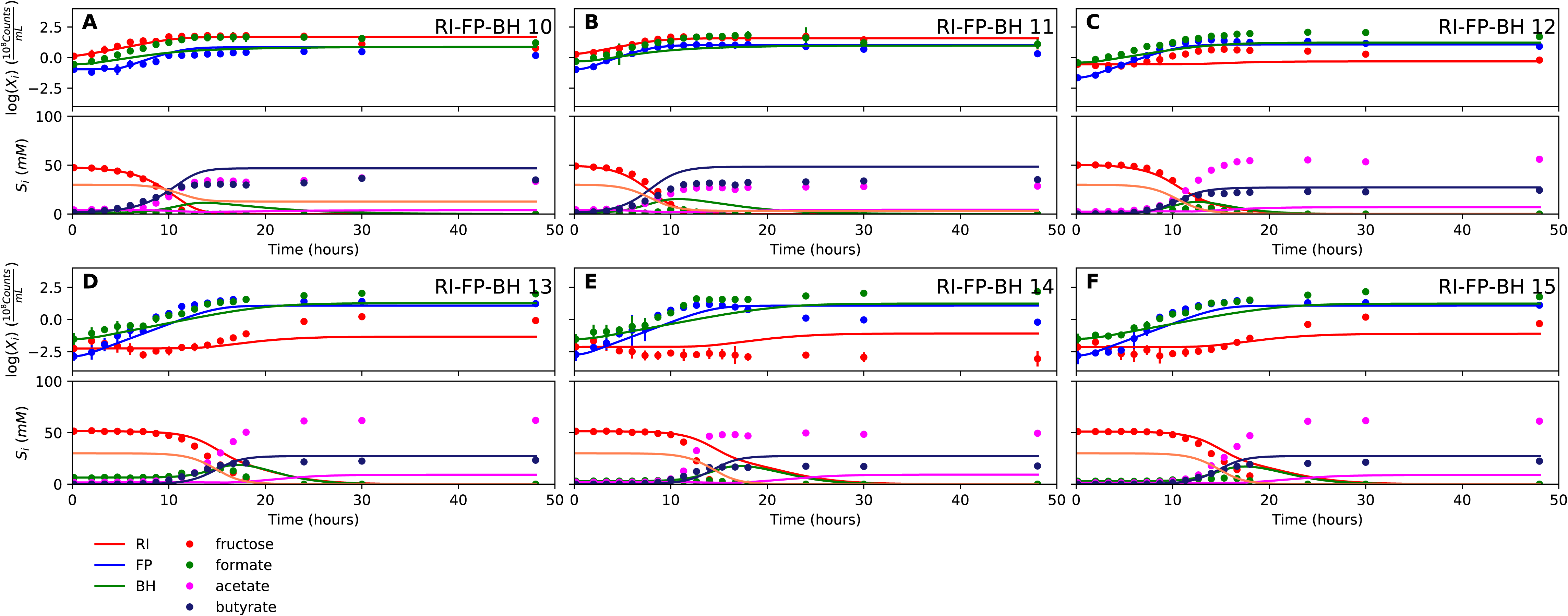
Fit to tri-culture experiments for the model parameterized on mono-cultures only. (A-B) Fit to tri-culture experiments dominated by *Roseburia intestinalis* and *Blautia hydrogenotrophica.* (C-F) Fit to tri-culture experiments dominated by *Faecalibacterium prausnitzii* and *Blautia hydrogenotrophica.* Lines represent model predictions and dots represent observations. The whiskers represent technical variation across triplicates. The shaded regions indicate the length of the estimated species-specific lag phases. The unknown compound represents an unspecified co-substrate assumed to be required by *Faecalibacterium prausnitzii.* Metabolites not included in the model are omitted from the plot. Experiment identifiers indicate which of the biological replicates is displayed. The model was parameterized on experiments RI_8, RI_14, FP_4, FP_15 and BH_14.

**Supplementary Figure 6.**
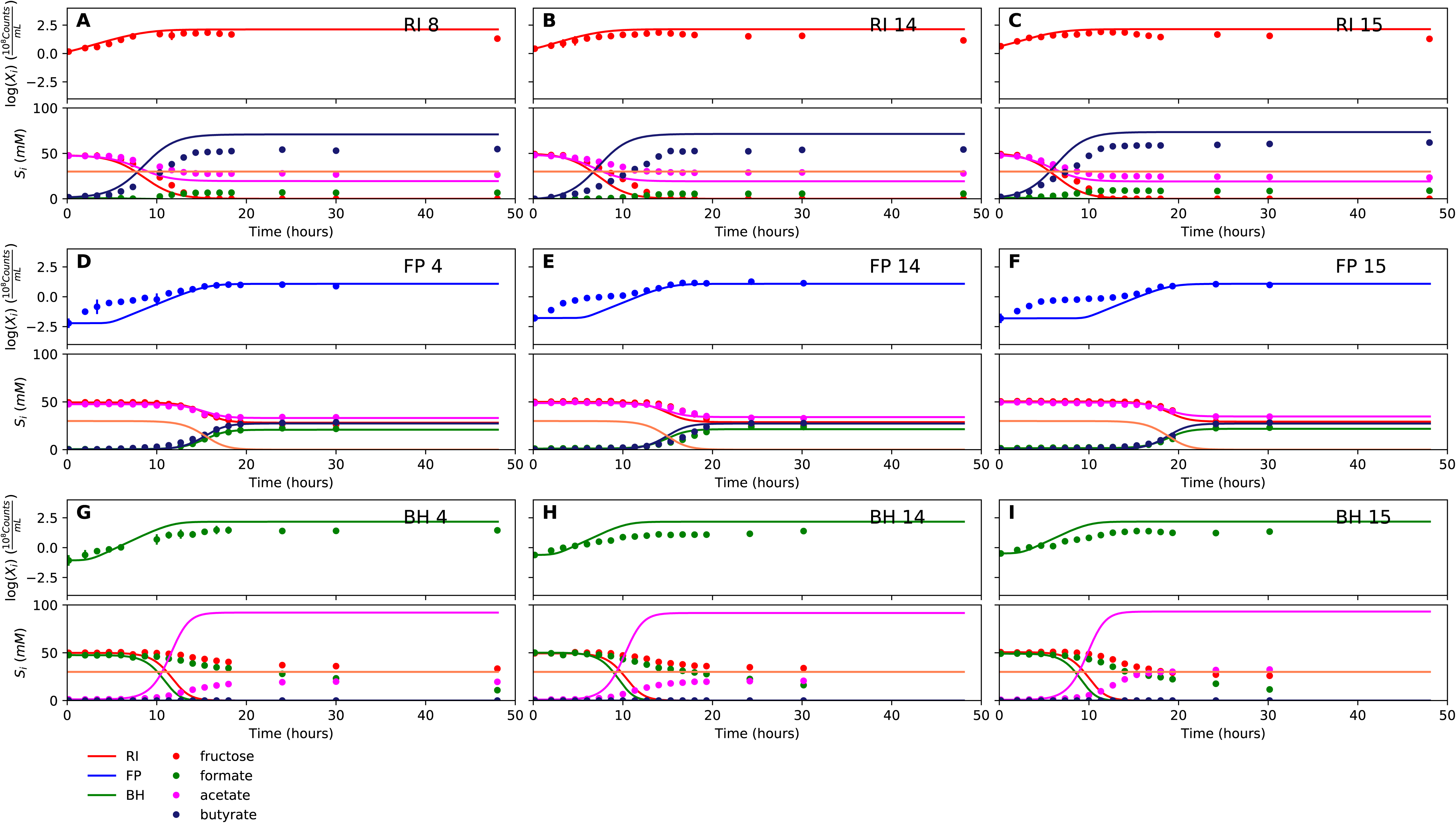
Fit to mono-culture experiments for the model parameterized on selected mono-cultures and bi-cultures. (A-C) Fit to *Roseburia intestinalis* mono-culture experiments. (D-F) Fit to *Faecalibacterium prausnitzii* mono-culture experiments. (G-I) Fit to *Blautia hydrogenotrophica* mono-culture experiments. Lines represent model predictions and dots represent observations. The whiskers represent technical variation across triplicates. The shaded regions indicate the length of the estimated species-specific lag phases. The unknown compound represents an unspecified co-substrate assumed to be required by *Faecalibacterium prausnitzii.* Metabolites not included in the model are omitted from the plot. Experiment identifiers indicate which of the biological replicates is displayed. The model was parameterized on experiments FP_4, FP_15, FP_BH_1, FP_BH_2 and RI_BH_4.

**Supplementary Figure 7.**
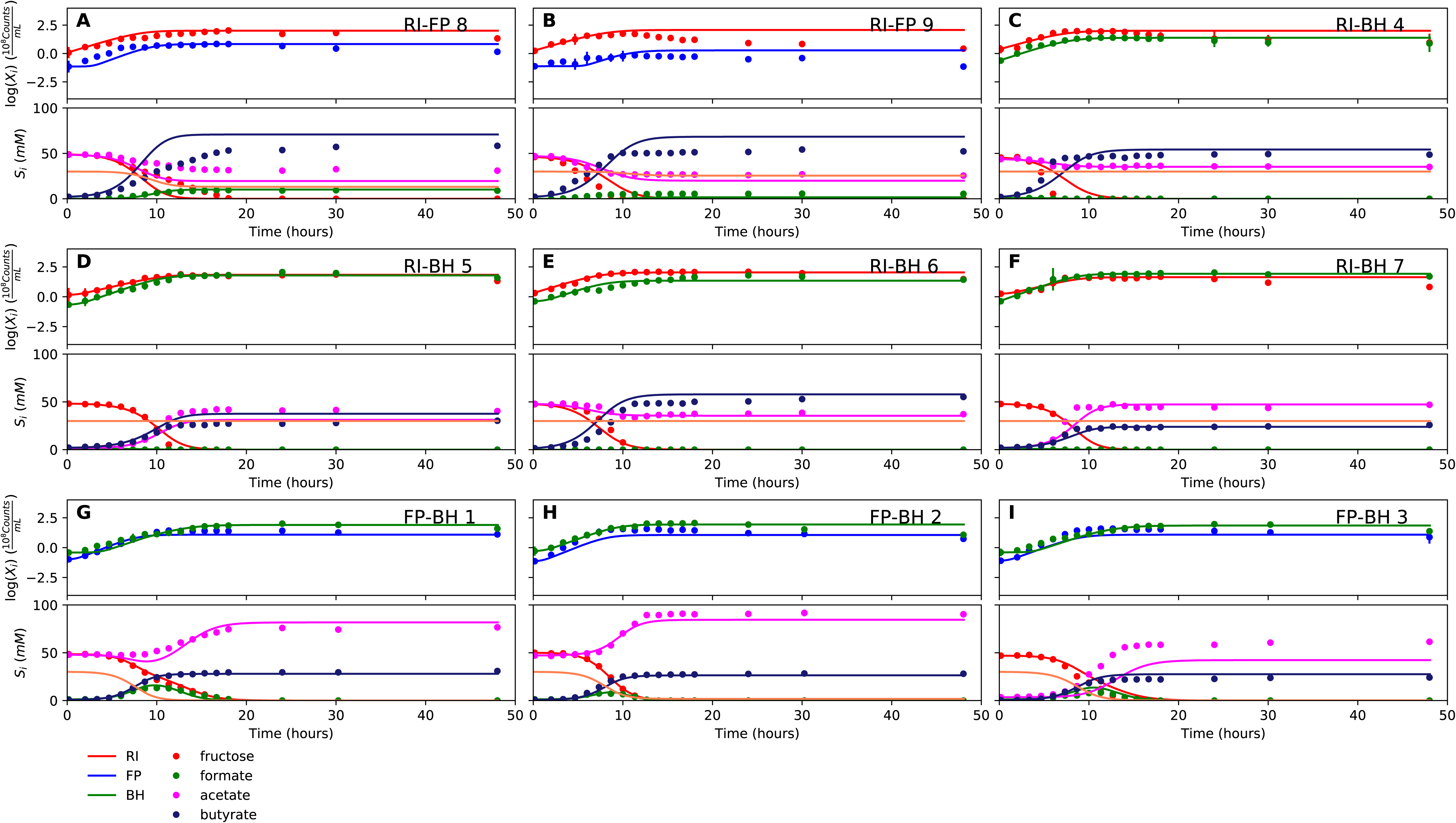
Fit to bi-culture experiments for the model parameterized on selected mono-cultures and bi-cultures. (A-B) Fit to *Roseburia intestinalis* and *Faecalibacterium prausnitzii* bi-culture experiments. (C-F) Fit to *Roseburia intestinalis* and *Blautia hydrogenotrophica* bi-culture experiments. (G-I) Fit to *Faecalibacterium prausnitzii* and *Blautia hydrogenotrophica* bi-culture experiments. Lines represent model predictions and dots represent observations. The shaded regions indicate the length of the estimated species-specific lag phases. The whiskers represent technical variation across triplicates. The unknown compound represents an unspecified co-substrate assumed to be required by *Faecalibacterium prausnitzii.* Metabolites not included in the model are omitted from the plot. Experiment identifiers indicate which of the biological replicates is displayed. The model was parameterized on experiments FP_4, FP_15, FP_BH_1, FP_BH_2 and RI_BH_4.

**Supplementary Figure 8.**
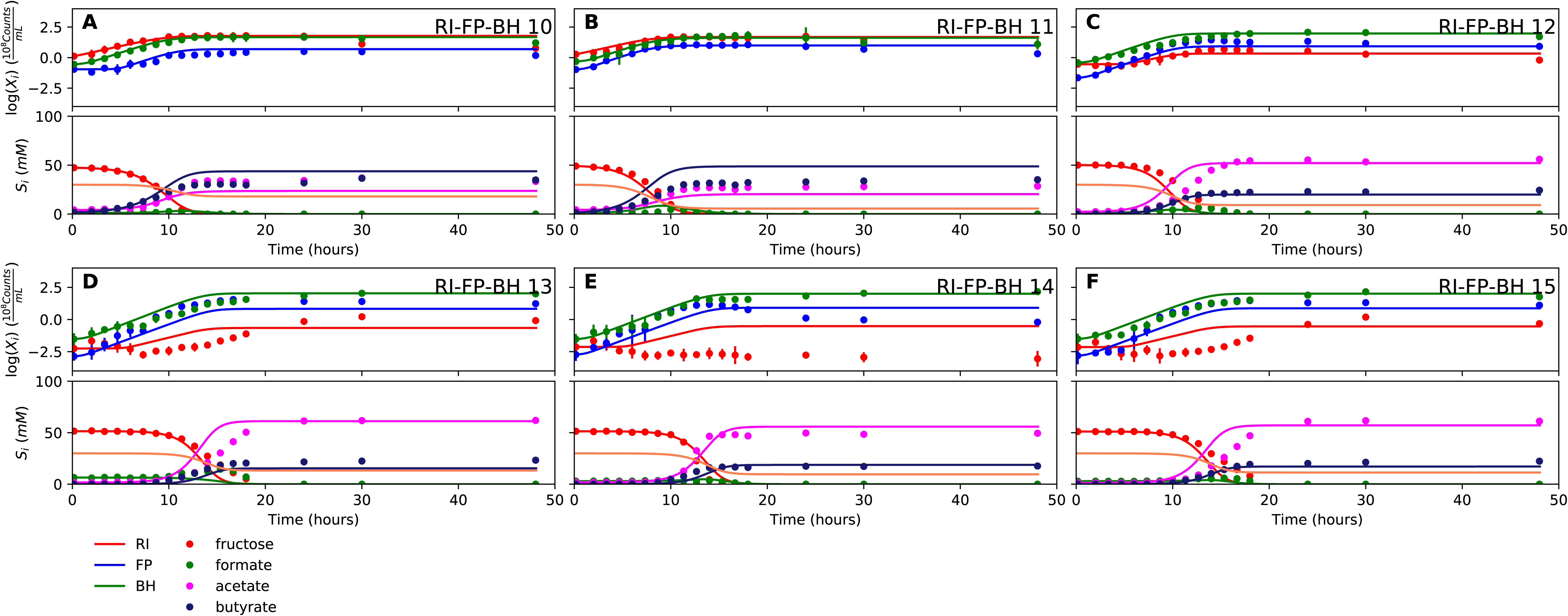
Fit to tri-culture experiments for the model parameterized on selected mono-cultures and bi-cultures. (A-B) Fit to tri-culture experiments dominated by *Roseburia intestinalis* and *Blautia hydrogenotrophica.* (C-F) Fit to tri-culture experiments dominated by *Faecalibacterium prausnitzii* and *Blautia hydrogenotrophica.* Lines represent model predictions and dots represent observations. The shaded regions indicate the length of the estimated species-specific lag phases. The whiskers represent technical variation across triplicates. The unknown compound represents an unspecified co-substrate assumed to be required by *Faecalibacterium prausnitzii.* Metabolites not included in the model are omitted from the plot. Experiment identifiers indicate which of the biological replicates is displayed. The model was parameterized on experiments FP_4, FP_15, FP_BH_1, FP_BH_2 and RI_BH_4.

**Supplementary Figure 9.**
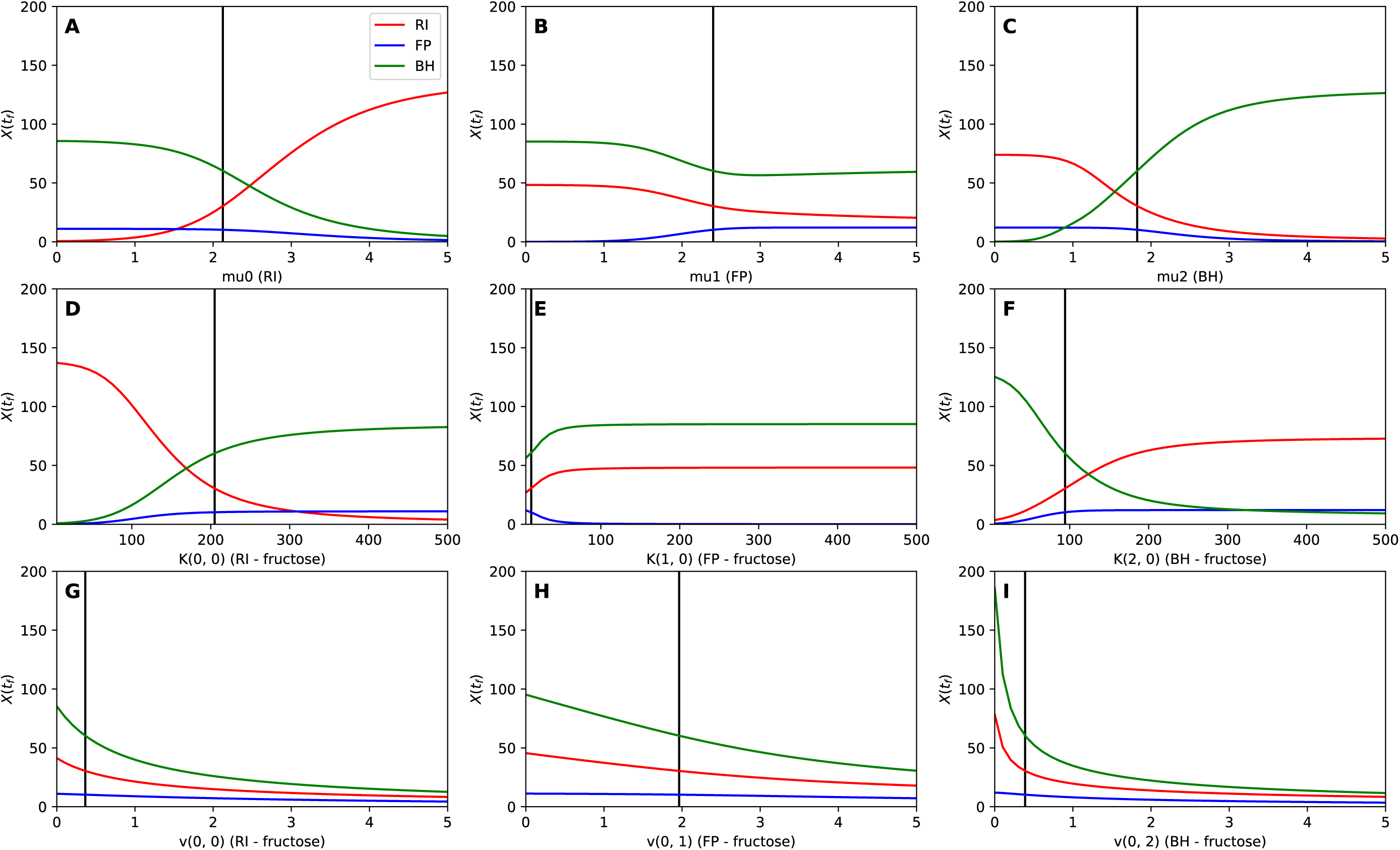
Sensitivity analysis. The abundance of *Roseburia intestinalis, Faecalibacterium prausnitzii* and *Blautia hydrogenotrophica* at the final time point is plotted, depending on the values of the maximal specific growth rate *μ* (A-C), the fructose half-saturation constant *K* (D-F) and the fructose uptake rate *ν* (G-I).

**Supplementary Table Legends**

Supplementary Table 1

A summary of all fermentation experiments is provided. Fermentation experiments selected for RNA-seq or model parameterization are marked accordingly.

Supplementary Table 2

The table provides parameter values after model parameterization on mono-cultures only and on mono- and bi-cultures (sheet 1), lag phases estimated from experiments (sheet 2) and predicted maximal abundances for the two parameterizations (sheet 3).

Supplementary Table 3

Table 3 lists genes that are significantly up- or down-regulated in tri-culture as compared to mono-culture for all three strains across all time points.

Supplementary Table 4

The sequences for species-specific primers and probes as well as their specificities are provided.

## References

Aharonovich D, Sher D (2016) Transcriptional response of Prochlorococcus to co-culture with a marine Alteromonas: differences between strains and the involvement of putative infochemicals. The ISME Jounal 10: 2892–2906

Anders S, Pyl PT, Huber W (2015) HTSeq—a Python framework to work with high-throughput sequencing data. Bioinformatics 31: 166–169

Andrews S (2010) FastQC: a quality control tool for high throughput sequence data. http://www.bioinformatics.babraham.ac.uk/projects/fastqc

Antharam VC, Li EC, Ishmael A, Sharma A, Mai V, Rand KH, Wang GP (2013) Intestinal Dysbiosis and Depletion of Butyrogenic Bacteria in Clostridium difficile Infection and Nosocomial Diarrhea. Journal of Clinical Microbiology 51: 28842892

Auchtung JM, Robinson CD, Britton RA (2015) Cultivation of stable, reproducible microbial communities from different fecal donors using minibioreactor arrays (MBRAs). Microbiome 3: 42

Baranyi J, Roberts T (1994) A dynamic approach to predicting bacterial growth in food. International Journal of Food Microbiology 23: 277–294

Bernalier A, Willems A, Leclerc M, Rochet V, Collins MD (1996) Ruminococcus hydrogenotrophicus sp. nov., a new H2/CO2-utilizing acetogenic bacterium isolated from human feces. Arch Microbiol 166: 176–183

Bolger AM, Lohse M, Usadel B (2014) Trimmomatic: a flexible trimmer for Illumina sequence data. Bioinformatics 30: 2114–2120

Buffie CG, Bucci V, Stein RR, McKenney PT, Ling L, Gobourne A, No D, Liu H, Kinnebrew M, Viale A, Littmann E, Brink MRMvd, Jenq RR, Taur Y, Sander C, Cross JR, Toussaint NC, Xavier JB, Pamer EG (2015) Precision microbiome reconstitution restores bile acid mediated resistance to Clostridium difficile. Nature 517: 205–208

Chiu H-C, Levy R, Borenstein E (2014) Emergent Biosynthetic Capacity in Simple Microbial Communities. PLoS Computational Biology 10: e1003695

Cremer J, Arnoldini M, Hwa T (2017) Effect of water flow and chemical environment on microbiota growth and composition in the human colon. PNAS 114: 6438–6443

D’hoe K, Conterno L, Fava F, Falony G, Vieira-Silva S, Vermeiren J, Tuohy K, Raes J (2018) Prebiotic Wheat Bran Fractions Induce Specific Microbiota Changes. Frontiers in Microbiology 9: 00031

de Vos MGJ, Zagorski M, McNally A, Bollenbach T (2017) Interaction networks, ecological stability, and collective antibiotic tolerance in polymicrobial infections. PNAS Early Edition

Degnan PH, Taga ME, Goodman AL (2014) Vitamin B12 as a modulator of gut microbial ecology. Cell Metab 20: 769–778

Duncan SH, Hold GL, Barcenilla A, Stewart CS, Flint HJ (2002a) Roseburia intestinalis sp. nov., a novel saccharolytic, butyrate-producing bacterium from human faeces. Int J Mol Evol Microbiol 52: 1615–1620

Duncan SH, Hold GL, Harmsen HJM, Stewart CS, Flint HJ (2002b) Growth requirements and fermentation products of Fusobacterium prausnitzii, and a proposal to reclassify it as Faecalibacterium prausnitzii gen. nov., comb. nov.. Int J Mol Evol Microbiol 52: 2141–2146

Falony G, Calmeyn T, Leroy F, Vuyst LD (2009a) Coculture Fermentations of Bifidobacterium Species and Bacteroides thetaiotaomicron Reveal a Mechanistic Insight into the Prebiotic Effect of Inulin-Type Fructans. Applied and Environmental Microbiology 75: 2312–2319

Falony G, Lazidou K, Verschaeren A, Weckx S, Maes D, Vuyst LD (2009b) In vitro kinetic analysis of fermentation of prebiotic inulin-type fructans by Bifidobacterium species reveals four different phenotypes. Applied and Environmental Microbiology 75: 454–461

Falony G, Verschaeren A, Bruycker FD, Preter VD, Verbeke K, Leroy F, Vuyst LD (2009c) In Vitro Kinetics of Prebiotic Inulin-Type Fructan Fermentation by Butyrate-Producing Colon Bacteria: Implementation of Online Gas Chromatography for Quantitative Analysis of Carbon Dioxide and Hydrogen Gas Production. Applied and Environmental Microbiology 75: 5884–5892

Falony G, Vlachou A, Verbrugghe K, Vuyst LD (2006) Cross-Feeding between Bifidobacterium longum BB536 and Acetate-Converting, Butyrate-Producing Colon Bacteria during Growth on Oligofructose. Applied and Environmental Microbiology 72: 7835–7841

Falony* G, Joossens* M, Vieira-Silva* S, Wang* J, Darzi Y, Faust K, Kurilshikov A, Bonder MJ, Valles-Colomer M, Vandeputte D, Tito RY, Chaffron S, Rymenans L, Verspecht C, Sutter LD, Lima-Mendez G, D’hoe K, Jonckheere K, Homola D, Garcia R et al. (2016) Population-level analysis of gut microbiome variation. Science 352: 560–564

Freilich S, Zarecki R, Eilam O, Segal ES, Henry CS, Kupiec M, Gophna U, Sharan R, Ruppin1 E (2011) Competitive and cooperative metabolic interactions in bacterial communities. Nature Communications 2: 589

Friedman J, Higgins LM, Gore J (2017) Community structure follows simple assembly rules in microbial microcosms. Nature Ecology and Evolution 1: 0109

Gause GF (1932) EXPERIMENTAL STUDIES ON THE STRUGGLE FOR EXISTENCE. Journal of Experimental Biology 9: 389–402

Gause GF (1934) The Struggle for Existence. Williams & Wilkins Co., Baltimore

Geirnaert A, Calatayud M, Grootaert C, Laukens D, Devriese S, Smagghe G, Vos MD, Boon N, Wiele TVd (2016) Butyrate-producing bacteria supplemented in vitro to Crohn’s disease patient microbiota increased butyrate production and enhanced intestinal epithelial barrier integrity. Scientific Reports 7: 11450

Gibson TE, Bashan A, Cao H-T, Weiss ST, Liu Y-Y (2016) On the Origins and Control of Community Types in the Human Microbiome. PLoS Computational Biology 12: e1004688

Girish PS, Haunshi S, Vaithiyanathan S, Rajitha R, Ramakrishna C (2013) A rapid method for authentication of Buffalo (Bubalus bubalis) meat by Alkaline Lysis method of DNA extraction and species specific polymerase chain reaction. Journal of Food Science Technology 50: 141–146

Grivet J (2001) Nonlinear population dynamics in the chemostat. Computing in Science Engineering 3: 48–55

Heinken A, Khan MT, Paglia G, Rodionov DA, Harmsen HJM, Thiele I (2014) Functional Metabolic Map of Faecalibacterium prausnitzii, a Beneficial Human Gut Microbe. Journal of Bacteriology 196: 3289–3302

Hunter JD (2007) Matplotlib: A 2D graphics environment. Computing In Science & Engineering 9: 90–95

Jones E, Oliphant T, Peterson P (2001) SciPy: Open Source Scientific Tools for Python. http://www.scipy.org/

Kettle H, Louis P, Holtrop G, Duncan S, Flint H (2015) Modelling the emergent dynamics and major metabolites of the human colonic microbiota. Environmental Microbiology 17: 1615–1630

Kibbe WA (2007) OligoCalc: An online oligonucleotide properties calculator. Nucleic Acids Research 35: 43–46

Kim HJ, Huh D, Hamilton G, Ingber DE (2012) Human gut-on-a-chip inhabited by microbial flora that experiences intestinal peristalsis-like motions and flow. Lab on a Chip 12: 2165–2174

Kopylova E, Noé L, Touzet H (2012) SortMeRNA: fast and accurate filtering of ribosomal RNAs in metatranscriptomic data. Bioinformatics 28: 3211–3217

Kraal L, Abubucker S, Kota K, Fischbach MA, Mitreva M (2014) The Prevalence of Species and Strains in the Human Microbiome: A Resource for Experimental Efforts. PLoS ONE 9: e97279

Lahti L, Salojãrvi J, Salonen A, Scheffer M, Vos WMd (2014) Tipping elements in the human intestinal ecosystem. Nature Communications 5: 4344

Langmead B, Salzberg SL (2012) Fast gapped-read alignment with Bowtie 2. Nature Methods 9: 357–359

Love MI, Huber W, Anders S (2014) Moderated estimation of fold change and dispersion for RNA-seq data with DESeq2. Genome Biology 15: 550

Markowitz VM, Chen I-MA, Palaniappan K, Chu K, Szeto E, Grechkin Y, Ratner A, Jacob B, Huang J, Williams P, Huntemann M, Anderson I, Mavromatis K, Ivanova NN, Kyrpides NC (2012) IMG: the integrated microbial genomes database and comparative analysis system. Nucleic Acids Research 40: D115–D122

Marzorati M, Vanhoecke B, Ryck TD, Sadabad MS, Pinheiro I, Possemiers S, Abbeele PVd, Derycke L, Bracke M, Pieters J, Hennebel T, Harmsen HJ, Verstraete W, Wiele TVd (2014) The HMI™ module: a new tool to study the Host-Microbiota Interaction in the human gastrointestinal tract in vitro. BMC Microbiology 14: 133

Mo ML, Palsson BØ, Herrgård MJ (2009) Connecting extracellular metabolomic measurements to intracellular flux states in yeast. BMC Systems Biology 3: 37

Moens F, Lefeber T, De Vuyst L (2014) Oxidation of metabolites highlights the microbial interactions and role of Acetobacter pasteurianus during cocoa bean fermentation. Applied and Environmental Microbiology 80: 1848–1857

Moens F, Verce M, De Vuyst L (2017) Lactate- and acetate-based cross-feeding interactions between selected strains of lactobacilli, bifidobacteria and colon bacteria in the presence of inulin-type fructans. International Journal of Food Microbiology 241: 225–236

Moens F, Weckx S, Vuyst LD (2016) Bifidobacterial inulin-type fructan degradation capacity determines cross-feeding interactions between bifidobacteria and Faecalibacterium prausnitzii. International Journal of Food Microbiology 231: 76–85

Momeni B, Xie L, Shou W (2017) Lotka-Volterra pairwise modeling fails to capture diverse pairwise microbial interactions. eLIFE 6: e25051

Monod J (1950) La technique de culture continue, theorie et applications. Annales d’Institut Pasteur 79: 390–410

Muñoz-Tamayo R, Giger-Reverdin S, Sauvant D (2016) Mechanistic modelling of in vitro fermentation and methane production by rumen microbiota. Animal Feed Science and Technology 220: 1–21

Newton DF, Macfarlane S, Macfarlane GT (2013) Effects of Antibiotics on Bacterial Species Composition and Metabolic Activities in Chemostats Containing Defined Populations of Human Gut Microorganisms. Antimicrobial Agents and Chemotherapy 57: 2016–2025

Ondov B, Philippy A (2017) Mash Screen: what’s in my sequencing run. https://genomeinformatics.github.io/mash-screen/

Ondov BD, Treangen TJ, Melsted P, Mallonee AB, Bergman NH, Koren S, Phillippy AM (2016) Mash: fast genome and metagenome distance estimation using MinHash. Genome Biology 17: 132

Orth JD, Thiele I, Palsson BØ (2010) What is Flux Balance Analysis? Nature Biotechnology 28: 245–248

Plichta DR, Juncker AS, Bertalan M, Rettedal E, Gautier L, Varela E, Manichanh C, Fouqueray Cn, Levenez F, Nielsen T, Doré Jl, Machado AMD, Evgrafov MCRd, Hansen T, Jørgensen T, Bork P, Guarner F, Pedersen O, Metagenomics of the Human Intestinal Tract (MetaHIT) Consortium, Sommer MOA et al. (2016) Transcriptional interactions suggest niche segregation among microorganisms in the human gut. Nature Microbiology: 16152

Rey FE, Faith JJ, Bain J, Muehlbauer MJ, Stevens RD, Newgard CB, Gordon JI (2010) Dissecting the in Vivo Metabolic Potential of Two Human Gut Acetogens. Journal of Biological Chemistry 285: 22082–22090

Rivera-Chávez F, Zhang LF, Faber F, Lopez CA, Byndloss MX, Olsan EE, Xu G, Velazquez EM, Lebrilla CB, Winter SE, Bäumler AJ (2016) Depletion of butyrate-producing Clostridia from the gut microbiota drives an aerobic luminal expansion of Salmonella. Cell Host Microbe 19: 443–454

Rivière A, Selak M, Lantin D, Leroy F, De Vuyst L (2016) Bifidobacteria and butyrate-producing colon bacteria: importance and strategies for their stimulation in the human gut microbiota. Frontiers in Microbiology 7: 979

Rudbeck L, Dissing J (1998) Rapid, simple alkaline extraction of human genomic DNA from whole blood, buccal epithelial cells, semen and forensic stains for PCR. Biotechniques 25: 588–592

Schmidt J, Riedele C, Regestein L, Rausenberger J, Reichl U (2011) A novel concept combining experimental and mathematical analysis for the identification of unknown interspecies effects in a mixed culture. Biotechnol Bioeng 108: 1900-1911

Shah P, Fritz JV, Glaab E, Desai MS, Greenhalgh K, Frachet A, Niegowska M, Matthew Estes2 CJ, Seguin-Devaux C, Zenhausern F, Wilmes P (2016) A microfluidics-based in vitro model of the gastrointestinal human–microbe interface. Nature Communications: 11535

Sher D, Thompson JW, Kashtan N, Croal L, Chisholm SW (2011) Response of Prochlorococcus ecotypes to co-culture with diverse marine bacteria. The ISME Jounal 5: 1125–1132

Smith H, Waltman P (1995) The Theory of the Chemostat: Dynamics of Microbial Competition. Cambridge University Press,

Stein RR, Bucci V, Toussaint NC, Buffie CG, Raetsch G, Pamer EG, Sander C, Xavier JB (2013) Ecological Modeling from Time-Series Inference: Insight into Dynamics and Stability of Intestinal Microbiota. PLoS Computational Biology 9: e1003388

T. Van de Wiele, P. Van den Abbeele, Ossieur W, Possemiers S, Marzorati M (2015) The Simulator of the Human Intestinal Microbial Ecosystem (SHIME®). In The Impact of Food Bioactives on Health, Verhoeckx K, Cotter P, López-Expósito I, Kleiveland C, Lea T, Mackie A, Requena T, Swiatecka D, Wichers H (eds) Springer, Cham

The UniProt Consortium (2017) UniProt: the universal protein knowledgebase. Nucleic Acids Research 45: D158–D169

Thiele I, Palsson BØ (2010) A protocol for generating a high-quality genomescale metabolic reconstruction. Nature Protocols 5: 93–121

Trosvik P, Rudi K, Næs T, Kohler A, Chan K-S, Jakobsen KS, Stenseth NC (2008) Characterizing mixed microbial population dynamics using time-series analysis. The ISME Jounal 2: 707–715

Trosvik P, Rudi K, Strætkvern KO, Jakobsen KS, Næs T, Stenseth NC (2010) Web of ecological interactions in an experimental gut microbiota. Environmental Microbiology 12: 2677–2687

Truong DT, Franzosa EA, Tickle TL, Scholz M, Weingart G, Pasolli E, Tett A, Huttenhower C, Segata N (2015) MetaPhlAn2 for enhanced metagenomic taxonomic profiling. Nature Methods 12: 902–903

Untergasser A, Nijveen H, Rao X, Bisseling T, Geurts R, Leunissen J (2007) Primer3Plus, an enhanced web interface to Primer3. Nucleic Acids Research 35: 71–74

Van Wey AS, Cookson AL, Roy NC, McNabb WC, Soboleva TK, Shorten PR (2014) Monoculture parameters successfully predict coculture growth kinetics of Bacteroides thetaiotaomicron and two Bifidobacterium strains. International Journal of Food Microbiology 191: 172–181

Venema K (2015) The TNO In Vitro Model of the Colon (TIM-2). In The Impact of Food Bioactives on Health, Verhoeckx K, Cotter P, López-Expósito I, Kleiveland C, Lea T, Mackie A, Requena T, Swiatecka D, Wichers H (eds) Springer, Cham

Wang H, Wei Z, Mei L, Gu J, Yin S, Faust K, Raes J, Deng Y, Wang Y, Shen Q, Yin S (2017) Combined use of network inference tools identifies ecologically meaningful bacterial associations in a paddy soil. Soil Biology and Biochemistry 105: 227–235

Wright ES, Vetsigian KH (2016) Inhibitory interactions promote frequent bistability among competing bacteria. Nature Communications 7: 11274

Ye J, Coulouris G, Zaretskaya I, Cutcutache I, Rozen S, Madden T (2012) Primer-BLAST: A tool to design target-specific primers for polymerase chain reaction. BMC Bioinformatics 13: 134

Zuker M (2003) Mfold web server for nucleic acid folding and hybridization prediction. Nucleic Acids Research 31: 3406–3415

